# *In situ* cryo-electron tomography of β-amyloid and tau in post-mortem Alzheimer’s disease brain

**DOI:** 10.1101/2023.07.17.549278

**Authors:** Madeleine A. G. Gilbert, Nayab Fatima, Joshua Jenkins, Thomas J. O’Sullivan, Andreas Schertel, Yehuda Halfon, Tjado H. J. Morrema, Mirjam Geibel, Sheena E. Radford, Jeroen J. M. Hoozemans, René A. W. Frank

## Abstract

A defining pathological feature of most neurodegenerative diseases is the assembly of proteins into amyloid that form disease-specific structures. In Alzheimer’s disease (AD) this is characterised by the deposition of amyloid-β (Aβ) and tau with AD-specific conformations. The *in situ* structure of amyloid in the human brain is unknown. Here, using cryogenic fluorescence microscopy (cryoFM)-targeted cryo-sectioning, cryo-focused ion beam scanning electron microscopy (cryoFIB-SEM) liftout and cryo-electron tomography (cryoET), we determined the in-tissue structure of β-amyloid and tau pathology in fresh post-mortem AD donor brain. β-amyloid plaques contained a mixture of fibrils and protofilaments arranged in parallel arrays and lattice-like structures, some of which were branched. Extracellular vesicles, extracellular droplets and open lipid bilayer sheets defined non-amyloid constituents of amyloid plaques. In contrast, tau inclusions were characterised by clusters of unbranched filaments. Subtomogram averaging of filaments within each cluster revealed distinct structures including variably twisted paired helical filaments (PHF) and chronic traumatic encephalopathy (CTE)-like tau filaments that were situated ∼1 μm apart within two microscopic regions of pathology. Filaments within a cluster were similar to each other, but different between clusters, showing that fibril heterogeneity is spatially organised and influenced by the subcellular tissue environment. The *in situ* structural approaches outlined here for targeting specific proteins within human donor tissues have applications to a broad range of neurodegenerative diseases.

## INTRODUCTION

Alzheimer’s disease is defined neuropathologically by the abnormal accumulation of aggregated Aβ peptides and tau that form extracellular and intracellular amyloid deposits, respectively^1^. Inherited forms of AD (familial AD; FAD) are caused by autosomal dominant mutations in the amyloid precursor protein (APP) and presenilin (PSEN1/PSEN2) genes^2, 3^. The APP gene encodes the precursor of Aβ, while the PSEN1/PSEN2 genes encode subunits of the ɣ-secretase complex that catalyse the final step of Aβ peptide production. Aβ peptides of varying lengths (including Aβ_1-40_, Aβ_1-42_ and Aβ_1-43_) are produced, of which Aβ_1-42_ is the major constituent of AD β-amyloid^4^. An immunotherapy raised against Aβ aggregates with the Arctic FAD mutation^5^, removes β-amyloid and delays the progression of AD^6^. Mutations in the microtubule associated protein tau (MAPT) gene cause neurodegeneration in the form of frontotemporal dementia and parkinsonism linked to chromosome 17, which result in tau pathology that lack β-amyloid deposits. In AD, the spread of aggregated tau correlates with neuronal loss and the sequence of cognitive decline^7, 8^.

Aβ and tau are highly aggregation prone, assembling into low-molecular weight oligomers or protofibrils that precede the formation of larger Aβ fibrils and tau filaments^9, 10^. Over decades, Aβ fibrils and tau filaments accumulate to form amyloid plaques and tau tangles in the parenchyma of the AD brain^11^. β-Amyloid plaques have been morphologically categorised as diffuse, dense-cored, fibrillar or neuritic^12, 13^, all of which contain Aβ_1-42_ fibrillar deposits^14, 15^. In addition, amyloid fibrils composed primarily of Aβ_1-40_ accumulate in and around blood vessels in various types of cerebral amyloid angiopathy^11, 16^. In contrast, tau filaments deposit within neuronal cell bodies and neurites forming tau tangles and tau threads, respectively^14, 15, 17^. At later stages, tau filaments can reside extracellularly in the form of ghost tangles and the remnants of atrophic neurites^14, 18, 19^.

The structure of Aβ_1-42_ fibrils purified from post-mortem AD brain and FAD mouse models was recently solved to high resolution by single particle cryoEM^20^. These *ex vivo* amyloids contain two structural conformers of Aβ_1-42_ amyloid (type I and type II), both of which are found in sporadic and familial AD cases^20^. These structures differ from Aβ_1-42_ fibrils prepared *in vitro*^21^ and from Aβ_1-40_ purified from the meninges of cerebral amyloid angiopathy (CAA) cases^22^. Atomic structures of tau filaments purified from AD^23, 24^ and other tauopathies^25–28^ suggest that tau forms disease-specific conformers. In AD, tau forms distinct ultrastructural polymorphs of paired helical filaments (PHF) and straight filaments (SF), both composed of three-repeat (3R) and four-repeat (4R) tau^23^. However, the molecular architecture and organization of Aβ and tau pathology within fresh, unfixed, human brain tissue remains unknown.

We recently reported the in-tissue molecular architecture of Aβ_1-42_ fibrils in a mouse model of FAD by cryogenic correlated light and electron microscopy (cryoCLEM) and cryo-electron tomography (cryoET) of tissue cryo-sections, identifying that these β-amyloid plaques are composed of fibrils, protofilament-like rods and branched amyloid, interdigitated by extracellular vesicles, extracellular droplets, and multilamellar bodies^29^. The extent to which this pathological architecture is representative of β-amyloid plaques in human AD brain is unknown. Additionally, FAD mouse models of β-amyloidosis do not recapitulate the full spectrum of Alzheimer’s disease pathology, including tau inclusions and neurodegeneration^30^.

Here, we determined the in-tissue three-dimensional architecture of β-amyloid plaques and tau pathology within fresh, hydrated, human post-mortem AD brain by cryoET. These data were collected using cryoFM to target specific pathology within cryo-sections and cryoFIB-SEM liftout lamella. Reconstructed tomographic volumes revealed extracellular β-amyloid plaques were composed of Aβ fibrils, branched fibrils and protofilament-like rods interlaced with non-amyloid constituents akin to our earlier studies of a murine FAD model^29^. Tau deposits consisted of unbranched filaments that formed clusters situated in intracellular and extracellular regions of the tissue. We determined the *in situ* structure of tau filaments to 13-18 Å resolution within each cluster by subtomogram averaging, which identified PHFs with variable filament twist, as well as chronic traumatic encephalopathy (CTE)-like tau filaments. Filaments within a cluster were similar to each other, but different between clusters, showing that fibril heterogeneity is spatially organised. The data suggest that tissue subcellular environments are key determinants of tau structure within each microscopic region of pathology.

## RESULTS

### Clinical history and neuropathology

CryoET was performed on post-mortem brain samples of the mid-temporal gyrus (GTM) from an AD donor and a non-demented donor (post-mortem delay 6:10 and 5:45 hours:min, respectively, **Methods**). The AD case was a 70-year-old woman with neuropathologically confirmed diagnosis following a 12-year history of progressive dementia. The donor started to have memory problems at the age of 54, and by the age of 58 was diagnosed with dementia, showing considerable memory deficits and disturbed executive functions. There was no family history of dementia and the genotype for APOE was e3/e3. Neuropathological analysis showed abundant amyloid plaques, tau tangles, tau threads, and very few CAA vessels across the GTM (**Fig. 1a** and **Extended Data Fig. 1a-d**). No α-synuclein, or TDP43 inclusions were detected, indicating the absence of pathologies that are associated with other neurodegenerative diseases (**Extended Data Fig. 1e-f**). To assess the AD donor tissue biochemically, sarkosyl insoluble tau aggregates were purified and immunoblotted (**Fig. 1b**). Tau aggregates were hyperphosphorylated and contained both 3R and 4R tau (**Extended Data Fig. 2a**). Overall, this immunohistochemical and biochemical profile is typical for AD cases^11^.

**Figure 1.**
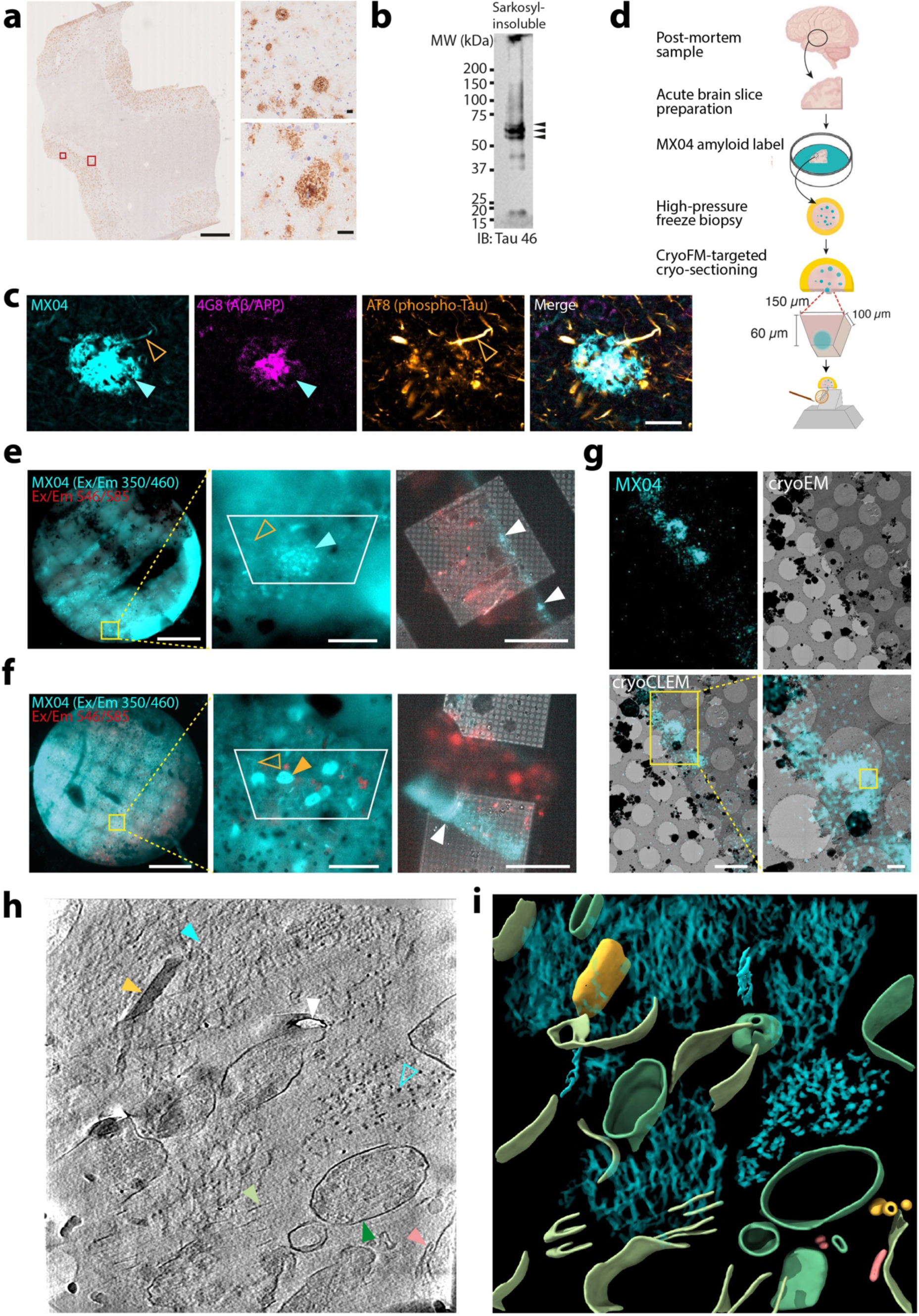
*In situ* cryoET of fresh AD post-mortem brain. **a-c** Neurohistopathological and biochemical characterisation of mid-temporal gyrus of AD post-mortem brain donor. **a** Immunohistochemical detection of Aβ/APP pathology (with 4G8, see **Methods**) in the grey matter of post-mortem AD donor. Scale bar, 2 mm. Large and small red rectangle, indicate close-up images shown in upper-and lower-right panels (scale bar, 20 μm), respectively. See also **Extended Data Fig. 1**. **b** Immunoblot detection of sarkosyl-insoluble tau probed with tau 46. Arrowheads, indicate full-length phospho-tau bands. See also **Extended Data Fig. 2a**. **c** Fluorescence confocal microscopic detection from left to right, amyloid (MX04), Aβ/APP (4G8), phospho-Tau (AT8), and merged in AD post-mortem donor brain. Cyan arrowhead, β-amyloid plaque. Open orange arrowhead, tau thread (containing hyperphosphorylated tau and amyloid indicated by MX04). Scale bar, 20 μm. See also **Extended Data Fig. 2b**. **d** Schematic showing workflow for the preparation of fresh AD post-mortem brain for *in situ* structure determination by cryoCLEM and cryoET. **e** CryoFM of high-pressure frozen AD post-mortem brain biopsy. Cyan, MX04 fluorescence. Red, autofluorescence detected with excitation and emission of 546 nm and 585 nm respectively. Left panel, high-pressure frozen fresh AD post-mortem brain biopsy. Scale bar, 0.5 mm. Middle panel, close-up. Cyan arrowhead, putative β-amyloid plaque. Open orange arrowhead, putative tau thread. White trapezium, area encompassing tissue stub that was subsequently trimmed and from which tissue cryo-sections were collected. Scale bar, 50 µm. Right panel, cryoFM of 70 nm thick tissue cryosection attached to EM grid that contained β-amyloid plaque. White arrowhead, MX04-labelled amyloid pathology. Scale bar, 50 µm. **f** Same as **e** but showing tau tangle and threads indicated by closed and open orange arrowheads, respectively. CryoET shown in Fig 2. **g** CryoCLEM targeting of MX04-labelled β-amyloid plaque in tissue cryo-section. Top left panel, cryoFM detection of MX04-labelled amyloid. Top right panel, cryoEM of MX04-labelled β-amyloid plaque cryo-section. Lower left panel, aligned images of cryoFM and cryoEM. Yellow rectangle, area shown as close up. Scale bar, 5 µm. Lower right, panel, close-up. Yellow rectangle, location from which cryoET data were collected (see **Extended Data Fig. 4a**). Scale bar, 1 µm. **h** Tomographic slice of β-amyloid pathology in AD post-mortem brain cryo-section (see **Extended Data Movie 1** and cryoET of non-demented donor control in **Extended Data Fig. 5**). Filled and open cyan arrowhead, fibril in the *x-y* plane and axially (*z*-axis) of the tomogram, respectively. Yellow arrowhead, extracellular droplet. Pink arrowhead, extracellular vesicle. Dark green arrowhead, subcellular compartment. Light green arrowhead, open membrane sheet. White arrowhead, knife damage. Scale bar, 10 nm. **i** Segmentation of tomogram coloured as in **h**.

**Figure 2.**
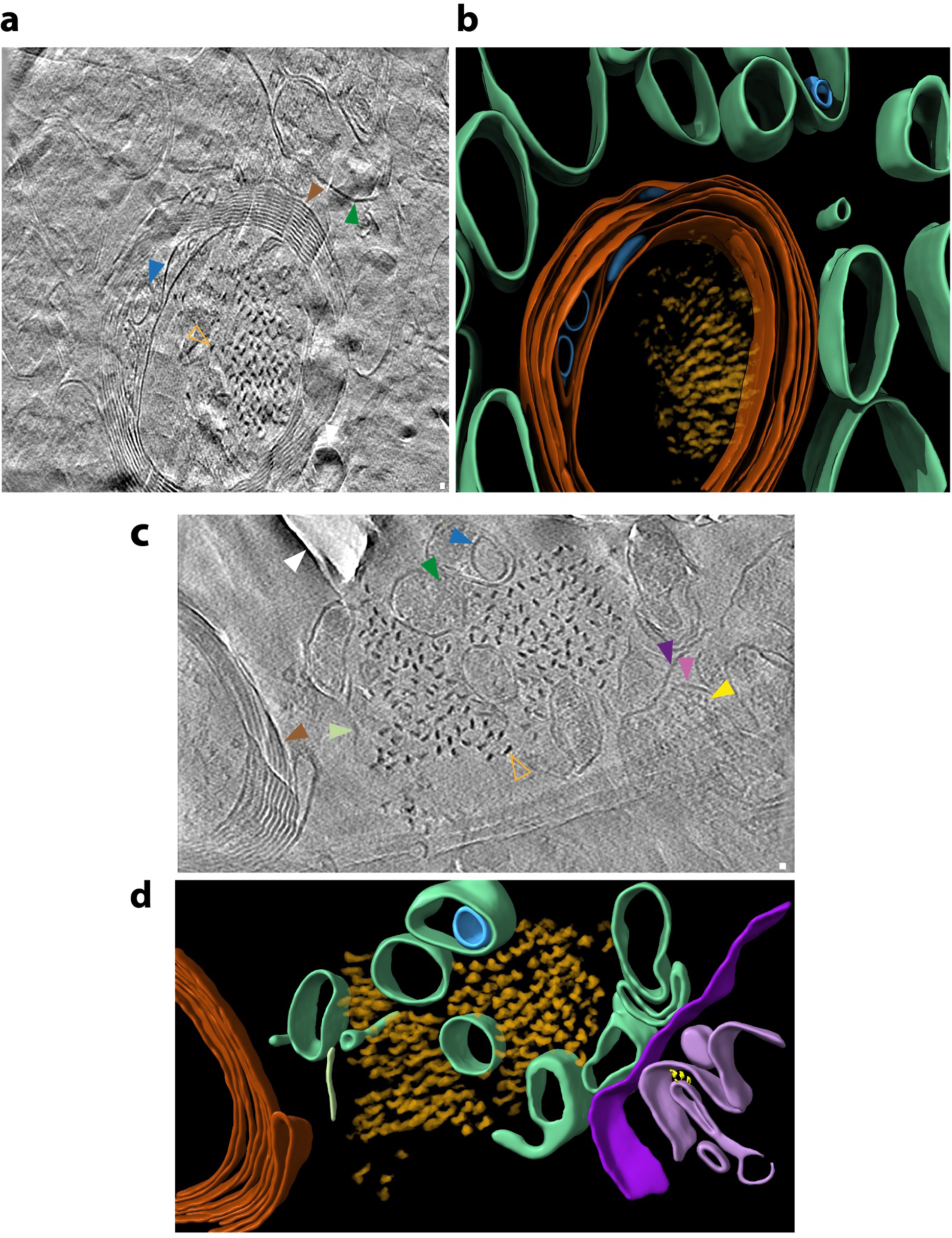
*In situ* cryoET of tau deposits in fresh AD post-mortem brain. **a** Tomographic slice of intracellular tau pathology in AD post-mortem brain cryo-section (see **Extended Data Movie 4** and **Extended Data Fig. 6**). Open orange arrowhead indicates filament oriented axially (*z*-axis) within the tomogram. Brown arrowhead indicates myelinated axon. Green arrowhead indicates sub-cellular compartment. Blue arrowhead, intracellular membrane-bound organelle. Scale bar, 10 nm **b** Segmentation of tomogram coloured as in **a**. **c** Tomographic slice of extracellular tau pathology in AD post-mortem brain cryo-section (see **Extended Data Movie 5**). Open orange arrowhead indicates tau filament oriented axially (*z*-axis) within the tomogram. Brown arrowhead indicates myelinated axon. Dark and light purple arrowheads indicate outer and inner membranes of damaged mitochondrion. Yellow arrowhead indicates putative Fo-F1 ATPase. Dark green arrowhead, subcellular compartment. Light green arrowhead, open membrane sheet. Blue arrowhead, intracellular membrane-bound organelle. White arrowhead, knife damage. Scale bar, 10 nm. **d** Segmentation of tomogram coloured as in **c**.

To detect amyloid deposits in fresh AD tissue, acute brain slices were incubated in methoxy-X04 (MX04), a generic amyloid fluorescent label^31^. Confocal microscopy showed wide-spread distribution of MX04-labelled amyloid, including 30-50 μm diameter deposits characteristic of β-amyloid plaques, as well as tau tangles and tau-filled neuropil threads (**Fig. 1c** and **Extended Data Fig. 2b**)^32^. Immunofluorescence detection of Aβ/APP and phospho-tau confirmed the identity of these morphologically distinct MX04-labelled deposits (**Fig. 1c** and Extended Data Fig.2b).

### *In situ* cryoCLEM of post-mortem brain

To locate β-amyloid and tau pathology within fresh post-mortem tissue, we adapted a workflow (**Fig. 1d**) from our earlier cryoET studies of brain tissue from healthy^33^ and FAD model^29^ mice. Acute brain slices of post-mortem AD donor tissue labelled with MX04, were vitrified by high-pressure freezing. Cryo-fluorescence microscopy (cryoFM) of these vitrified tissues revealed the location of MX04-labelled amyloid pathology, including neuritic plaques (**Fig. 1e**), tau tangles (**Fig. 1f**) and threads (**Fig. 1e-f**), which resembled those in fixed tissue (**Fig. 1c** and **Extended Data Fig. 2b**). These pathologies were absent in MX04-labelled non-demented control tissue (**Extended Data Fig. 2c**).

To prepare vitrified post-mortem brain for cryoEM, 70 nm thick tissue cryo-sections were collected (**Fig. 1d-f**) from a MX04-labelled β-amyloid plaque (**Fig. 1e**) and from a second location enriched in tau tangles and threads (**Fig. 1f** and **Extended Data Fig. 2d**). CryoFM confirmed the presence of MX04-labelled amyloid within these tissue cryo-sections (**Fig. 1e-f**). MX04-labelled amyloid was mapped by cryoCLEM onto medium magnification electron micrographs (**Fig. 1g**). In regions with strong MX04 signal, putative amyloid was directly observed by cryoEM (**Extended Data Fig. 3a**).

**Figure 3.**
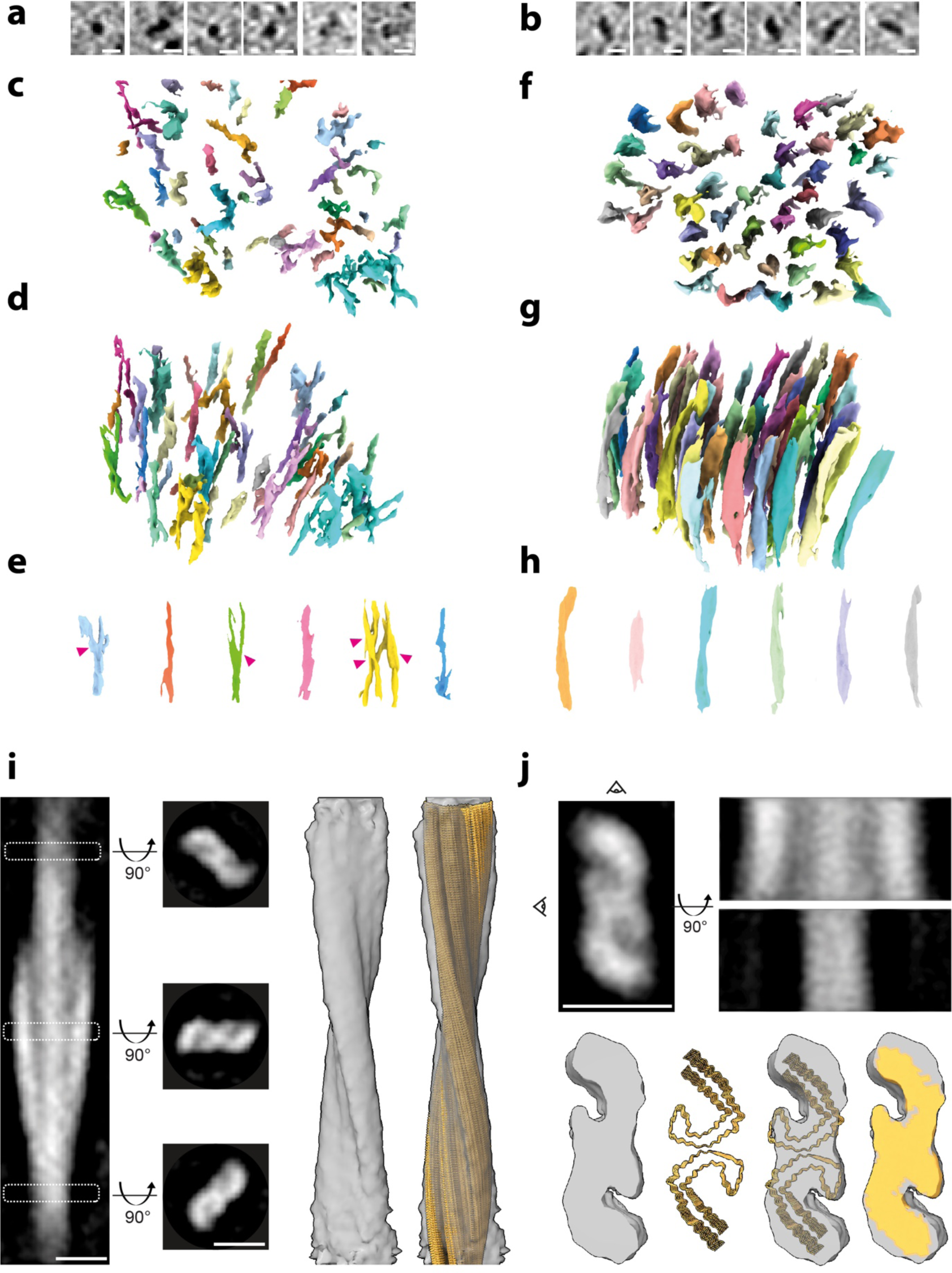
In-tissue architecture of β-amyloid fibrils and tau filaments, and subtomogram averaging of tau in fresh post-mortem brain cryo-sections. **a** Tomographic slices showing panel of β-amyloid fibrils oriented axially (*z*-axis) within the tomogram. Scale bar, 10 nm. **b** Tomographic slices showing a panel of tau filaments oriented axially (*z*-axis) within the tomogram. Scale bar, 10 nm. **c** and **d** show top and side view of raw tomographic density containing a lattice of β-amyloid, respectively. Each individual amyloid fibril is coloured differently. 33% (17 out of a total of 51) individually coloured fibrils had branch points. **e** Side views of raw tomographic density of single amyloid fibrils from within the lattice/cluster. Magenta arrowhead, branch points. **f**, **g** and **h** show the same as **c**, **d** and **e**, but for tau. **i** Subtomogram average of 121 tau filaments (stalkInit, see **Methods**) located extracellularly (from one tomogram, see **Fig. 2c-d** and **Extended Data Movie 5**). Left panel, side view of tomographic slice through averaged volume showing helical twist. White dashed rectangles, position of middle left panels along filament axis of three top view tomographic slices at three positions (25 voxels, 23.75 nm apart) showing a pair of C-shaped protofilaments consistent with the substructure of *ex vivo* purified tau PHF^23^. Middle right and right panels, subtomogram average map of tau filament with and without helically averaged atomic model of tau PHF (PDB 5osl) composed of two protofilaments (coloured yellow)^23^ fitted into the map. Scale bars, 10 nm. **j** Filament sub-volume (addModPts, n=950, see **Methods**) average of tau amyloid from one AD cryo-section tomogram (see Fig. 2 and **Extended Data Movie 5**) (12.9 Å resolution at FSC 0.5, see **Extended Data Fig. 9**). Top left, middle and right panels, tomographic slices through averaged map from top and two side views rotated 90° about the filament axis indicated by eye-symbols, respectively. Lower left, top view of averaged map. Lower middle left and right, helically averaged atomic model of tau PHF (PDB 5osl)^23^ alone and fitted into averaged map, respectively. Each protofilament in the PHF is coloured yellow. Lower right, subtomogram map is shown coloured by the occupancy of the fitted atomic model showing remaining regions of the map unaccounted for by the model in grey. Scale bars, 10 nm.

CryoCLEM maps were used to target the collection of tomographic tilt series, each encompassing a ∼1 μm^2^ area of the tissue cryosection (2.4 Å pixel size). We collected 42 and 25 tomograms in and around regions of cryo-sections that contained an MX04-labelled β-amyloid plaque (**Extended Data Table 1**) and tau tangles (**Extended Data Table 2**), respectively. An additional 64 tomograms collected from non-demented donor tissue cryo-sections were used as a control (**Extended Data Table 3**). Reconstructed tomographic volumes revealed the native, in-tissue, three-dimensional molecular architecture of AD pathology in fresh post-mortem brain (**Fig. 1h-i** and **Fig. 2**, see also **Extended Data Movies 1-3** from amyloid plaques**, Extended Data Movies 4-6** from tau tangles and threads, and **Extended Data Movies 7-9** from non-demented control).

### In-tissue architecture of a β-amyloid plaque and tau inclusions

Fibrils were apparent in all tomographic volumes collected from the MX04-labelled β-amyloid plaque (**Extended Data Table 1**). No fibrils were present in any of the control donor tomograms (n=64, **Extended Data Table 3**). The fibrils within β-amyloid plaques formed parallel arrays or a lattice (**Fig. 3c-d**, **Fig. 1h-i and Extended Data Fig. 4**). This architecture is similar to that we reported in the amyloid plaques of an FAD mouse model^29^. In a subset of tomograms, the extracellular location of the amyloid fibrils could be determined unambiguously by its juxtaposition to subcellular compartments enclosing intracellular organelles or a myelinated axon (**Extended Data Fig. 4 and Extended Data Table 1**).

**Figure 4.**
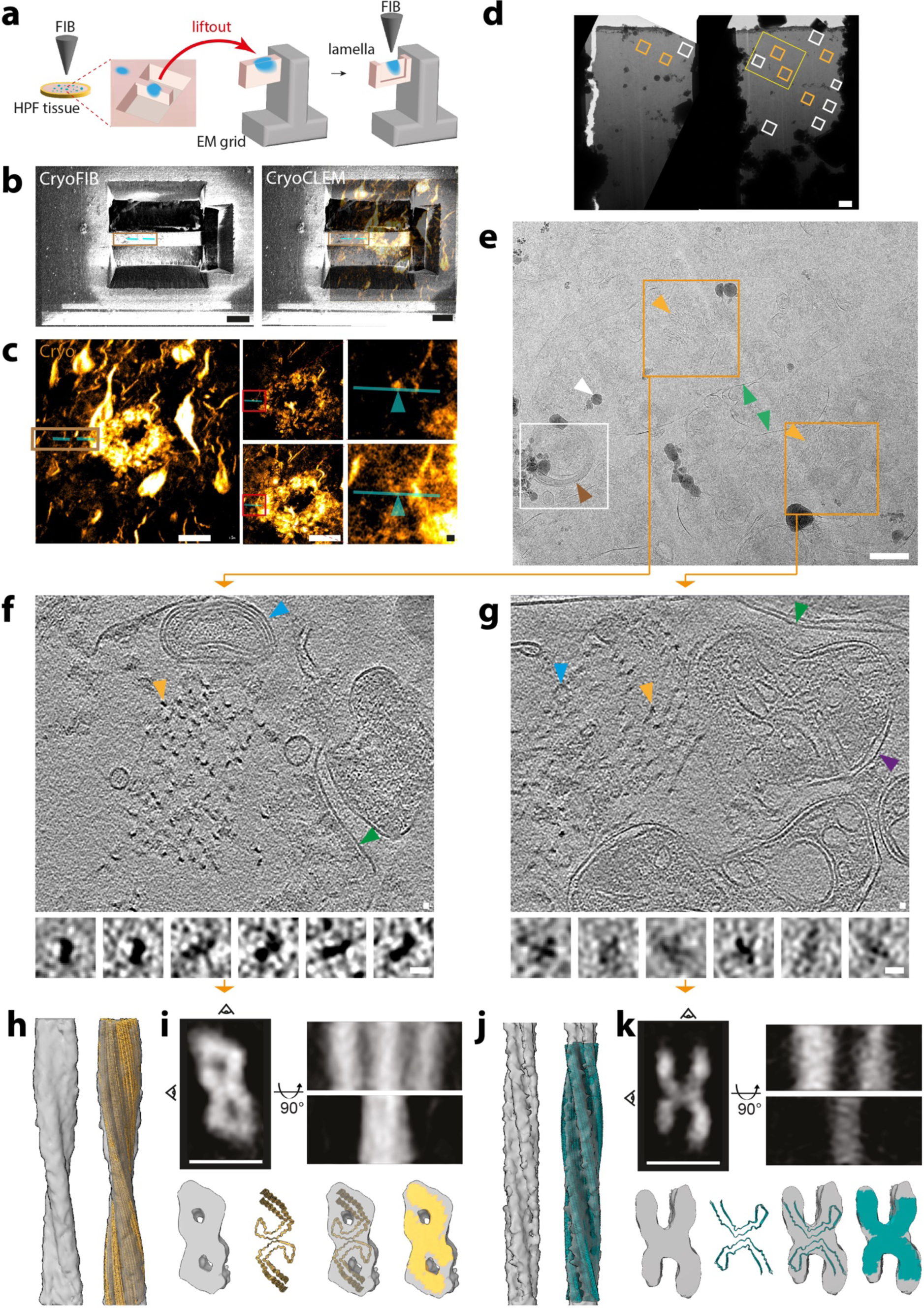
cryoCLEM-targeted cryoFIB-SEM liftout of tau thread lamellae from post-mortem AD brain. **a** Schematic summarising cryoCLEM-targeted cryoFIB-SEM liftout workflow. High pressure frozen tissue containing MX04-labelled amyloid was focused ion beam (FIB) milled to isolate an 80 μm wide, 10 μm thick, ∼30 μm deep tissue chunk containing a MX04-labelled tau (blue). The chunk was lifted out and attached to a half-moon cryoEM grid. The tissue chunk was then thinned to produce 130-200 nm thick tissue lamella windows. **b** Left, cryoFIB image of HPF frozen brain showing normal view of milled tissue chunk that preceded liftout and cryoFIB-SEM thinning of tissue lamella. Right, FIB image aligned with confocal cryoFM of MX04 amyloid in fresh high-pressure frozen tissue. Brown rectangle, corresponds to proximal tissue chunk that was lifted out. Cyan lines, correspond to the locations of the tissue lamella windows within the tissue chunk that was FIB-milled after liftout. See **Extended Data Fig. 12 and 13** and **Methods**). Scale bar, 20 μm. **c** Three-dimensional (*z*-stack) super-resolution cryoFM imaging of MX04 labelled amyloid in post-mortem AD brain tissue targeted for the preparation of cryoFIB-SEM liftout lamella. Left, extended depth of field image of confocal *z*-stack. Middle, super-resolution cryoFM images showing top and bottom of the tissue corresponding to the *z*-slices above and below (1.9 μm apart), respectively. Scale bar 20 μm. Red rectangle, regions shown enlarged in right panels. Cyan line, show location of tissue lamella window. Cyan arrowhead, microscopic regions of MX04-labelled amyloid corresponding to the locations above and below of the first and second tomograms that were collected (see panels **f** and **g**), respectively. Scale bar, 1 μm. See also **Extended Data Figs. 12-14.** **d** Overview electron micrograph of cryoFIB-SEM liftout with two lamella windows. Orange rectangle, location of tomogram collected that contained tau filaments. White rectangle, location of tomograms that lacked tau filaments. Yellow rectangle, region enlarged in **e**. Scale bar, 1 μm. **e** Close up of **d**. Orange rectangles, regions of tissue lamella containing tau clusters (orange arrowhead) within two membrane-bound subcellular compartments (green arrowheads). White arrowhead. ice contaminations. White rectangle, location of tomogram containing myelinated axon (brown arrowhead) and lacking tau filaments (see **Extended Data Movie 12**). Scale bar, 500 nm. **f** and **g** Top panel, tomographic slices from tissue lamella of tau thread. Orange arrowhead, single tau filament. Purple arrowhead, mitochondrion. Green arrowhead, membrane enclosing subcellular compartment. Blue arrowhead, intracellular membrane-bound organelle. Scale bar, 10 nm (see **Extended Data Movies 10 and 11**). Bottom panels, tomographic slices showing representative cross section of filaments. Scale bar, 10 nm. **h** Subtomogram average of 68 tau filaments (StalkInit, see **Methods**). Left and right, subtomogram average map and helically averaged atomic model of tau PHF (each filament in the PHF coloured yellow) (PDB 5osl)^23^ fitted into the subtomogram average map. **i** Subtomogram average of tau filament sub-volumes (addModPts, n=900, 15.4 Å resolution at FSC 0.5 see **Methods** and **Extended Data Fig. 15b**). Top left, middle and right panels, tomographic slices through averaged map from top and two side views rotated 90° about the filament axis indicated by eye-symbols, respectively. Lower left, top view of averaged map. Lower middle left and right, atomic model of helically averaged tau PHF (yellow, PDB 5osl)^23^ and fitted into subtomogram average map, respectively. Lower right, subtomogram map is shown coloured by the occupancy of the fitted atomic model (yellow). Scale bar, 10 nm. **j-k** Same as **j-i** but for second cluster of tau filaments located ∼1 μm away (62 filaments from which 1258 sub-volumes were averaged to 13.4 Å resolution at FSC 0.5, see **Extended Data Fig. 15c** and **Methods**) and with atomic model of tau CTE type 1 (cyan, PDB 6nwp)^46^ fitted into the subtomogram average.

β-amyloid plaques were interdigitated by various membrane-bound subcellular compartments and associated macromolecular constituents (**Fig. 1h-i, Extended Data Fig. 4 and Extended Data Table 1**). These included a prevalence of extracellular vesicles, extracellular droplets, and open membrane sheets. All except open membrane sheets were comparable with those we observed by cryoET in an FAD mouse model^29^. These non-amyloid constituents were absent from non-demented donor control tissue tomograms (**Extended Data Fig. 5 and Extended Data Table 3**).

Tomograms collected from MX04-labelled tau-containing tissue cryo-sections revealed filaments arranged as a 300-800 nm parallel cluster (**Fig. 2e, Extended Data Fig. 6** and **Extended Data Table 2**), which were absent in control tissue cryo-sections (**Extended Data Table 3** and **Extended Data Fig. 5**). These deposits were within the cytoplasm of neurites and within myelinated axons (**Fig. 2a**). A cluster of this amyloid was also found that was not enclosed by a plasma membrane, but was located outside of a myelinated axon (**Fig. 2b**). Non-amyloid constituents of intracellular and extracellular MX04-labelled tau deposits included vesicles and damaged mitochondria that were in close proximity to tau clusters (**Fig. 2 and Extended Data Fig. 6**). These types of membrane-bound organelles were also present in non-demented control tissue cryosections in the absence of tau tangles (**Extended Data Table 3**).

To assess further the architecture and identity of fibrils in β-amyloid plaques and tau deposits, populations from each were compared. The cross section of fibrils from β-amyloid plaques and tau pathology-containing tomograms showed fibrils and filaments of 6 ± 2 nm and 15 ± 3 nm (mean ± SD) maximum diameter, respectively (**Fig. 3a-b** and **Extended Data Fig. 7a**). These widths were broadly consistent with the expected dimensions of atomic structures of *ex vivo* purified β-amyloid fibrils^20^ and tau filaments^23^, respectively, and confirm the MX04-targeted collection of tomographic volumes from β-amyloid plaques and tau inclusions. Amyloid plaque fibrils and tau filaments were oriented both in the *x*-*y* plane and axially in the *z*-direction of the tomographic volume. The latter provided sufficient contrast to trace individual fibrils throughout the tomographic volume (**Fig. 3**). In such regions of β-amyloid plaque tomograms, 33-40% of fibrils exhibited branch points, in which fibrils bifurcated from one another (**Fig. 3c-e**). This branched amyloid architecture was similar to that in an FAD mouse model of β-amyloidosis^29^. In tau tomograms, only unbranched, 8-20 nm thick filaments were present (**Fig. 3f-h**).

### Subtomogram averaging of in-tissue tau filaments and β-amyloid fibrils

To obtain higher resolution structural information we performed subtomogram averaging of amyloid in tau tangles and β-amyloid plaques. Since the subcellular environment within different locations of the tissue could influence the structure of the filament, we aligned and averaged subvolumes from each cluster of tau filaments independently. The averages obtained from seven distinct tau clusters showed filaments in which the helical twist was resolved (**Fig. 3i** and **Extended Data Fig. 8**). The highest resolution average (12.9 Å at 0.5 FSC, **Extended Data Fig. 9 & 16**) corresponded to an extracellular cluster of tau filaments, in which the C-shaped conformation of two protofilaments that compose PHFs was unambiguous (**Fig. 3j**). To compare this *in situ* structure with available helically averaged *ex vivo* structures of tau filaments^23–28^, we fitted atomic models in the subtomogram average map. The AD PHF model was best accommodated, whereas SF and all tau filament conformers from other tauopathies^23–28^ gave a poorer fit to the tomographic density (**Extended Data Fig. 9**). Subtomogram averages from five other tau clusters within the tissue also produced maps in which AD PHFs were best accommodated (**Extended Data Fig. 10a-e**). Only one tau cluster did not accommodate PHFs or SFs and higher resolution features were absent that could unambiguously indicate a specific tau amyloid fold (**Extended Data Fig. 10f**).

Comparing the subtomogram average helical parameters of each tau filament cluster with that of the *ex vivo* purified atomic model of PHF^23^ indicated that *in situ* tau filaments exhibited location-specific variability in helical twist (**Extended Data Figs. 7b** & **8**). All *in situ* PHF subtomogram averages were 17 to 74% less twisted than that of *ex vivo* PHFs (87-299 nm versus 71 nm cross over distance of *in situ* PHF-like filaments versus *ex vivo* PHF, respectively). The tau cluster that did not accommodate *ex vivo* atomic models of PHF or SF was 76% less twisted (**Extended Data Fig. 7b**). To test further the location specificity of this variation in tau filament structure, we pooled filament sub-volumes from all locations and performed unsupervised classification by principal component analysis (PCA). PCA classified the majority of filaments by subcellular location (**Extended Data Fig. 7c**), supporting that each tau cluster comprises structurally similar filaments and that all except one of the clusters were structurally different from one another.

In β-amyloid plaques, fibrils of varying width were apparent (**Fig. 3a and Extended Data Fig. 7a**). We therefore performed subtomogram averaging of all or sub-populations of amyloid deposits with different widths (protofilament-like rods, fibrils and thick fibrils), none of which resolved the helical twist or higher resolution structural features (**Extended Data Fig. 11**). This is not surprising since Aβ fibrils are smaller and less obviously featured structures^20^ than tau filaments^28^. Available atomic models of *ex vivo* purified Aβ type 2, type 1b^20^, and a single type 2 protofilament were nonetheless well accommodated into the averaged maps of fibrils, thick fibrils, and protofilament-like rods, respectively (**Extended Data Fig. 11**). This suggests that β-amyloid plaques contain a heterogeneous mixture of fibrillar species similar to that observed in a mouse model of AD^29^.

### cryoFIB-SEM liftout of post-mortem AD brain

The achievable resolution of subtomogram averaging in cryo-sections could be limited by knife damage to the sample that arises during sample preparation^34^. Therefore, we established an alternative workflow to prepare tissue lamella^35^ from fresh post-mortem AD brain by cryoFIB-SEM liftout (**Fig. 4a**). Three-dimensional super-resolution cryoFM was used to target the preparation of tissue lamellae to a location containing tau threads that were situated adjacent to a MXO4-labelled amyloid plaque (**Fig. 4b-c** and **Extended Data Figs. 12, 13 and 14).** Thirteen tomograms were collected throughout the lamella, six of which contained tau clusters (**Fig. 4d**).

Subtomogram averaging was performed with two tomograms that contained the highest copy number of tau filaments. These two tau clusters were separated by ∼1 μm within two membrane-bound subcellular compartments and were mapped to two distinct MX04-labelled tau threads located at the periphery of an MX04-labelled amyloid plaque (**Fig. 4e-g and Extended Data Figs. 12 & 13**). The cross-section of filaments appeared different within each tau cluster (**Fig. 4f-g**). Aligning filaments from each location independently produced structurally distinct subtomogram averages (**Fig. 4h-i** compared to **Fig. 4j-k**). The first cluster resembled PHFs from cryosections (94 nm crossover distance, **Extended Data Fig. 15a**), whereas the second cluster was structurally distinct and ∼2-fold more twisted (52 nm crossover distance).

Further averaging of smaller subvolumes along the helical axis of filaments in each cluster produced 13.4-15.4 Å resolution maps that revealed the conformation of protofilaments that compose tau filaments (**Fig. 4i** and **4k** and **Extended Data Fig. 15b-c, 17 & 18**). In the first cluster the atomic model of AD PHF^23, 24^ was well accommodated (**Fig. 4i**). The second cluster was structurally different with C-shaped protofilaments adopting a more open conformation compared to the first cluster (∼47 Å and ∼33 Å distance between the ends of each C-shaped protofilament, respectively). We fitted all available tau filament atomic models into the map. The CTE type-1 tau conformer^36^, associated with repetitive head impacts or concussion^37^, showed good overall agreement with the subtomogram averaged map (**Fig. 4j-k**). All other available structures of *ex vivo* purified tau filament conformers, including PHF and SF that are associated with AD^23^, did not fit in the tomographic density (**Extended Data Fig. 15b-c**). These *in situ* cryoET data indicate that tau filaments form clusters and highlights the co-existence of multiple distinct tau filament structures organised within two neighbouring microscopic regions of pathology.

## DISCUSSION

Light microscopic characterisation of amyloid in AD brain has formed the basis of diagnostic and disease classification over decades. Recent atomic models of *ex vivo* AD Aβ fibrils and tau filaments prepared by bulk purification from whole brain regions have elucidated fibril and filament conformers specific to AD and other neurodegenerative diseases^20, 22, 28, 38^. Here, using cryoFM to guide cryoET and subtomogram averaging we have delineated the relationship between molecular structure, cellular context and the characteristic pattern of microscopic neuropathology in AD. These *in situ* structures of β-amyloid plaques and tau pathology revealed the heterogeneity of Aβ fibrils and the spatial segregation of distinct tau filament structures within different cellular contexts from a single brain region of an individual post-mortem AD donor.

β-amyloid plaques were characterised by a lattice-like architecture of amyloid fibrils interdigitated by non-amyloid constituents, including extracellular vesicles and droplets, which were absent from non-demented post-mortem tissue tomograms. These were consistent with a recent cryoET study of a mouse model of AD^29^ and plastic-embedded EM of AD brain^14, 15^. We suggest these non-amyloid constituents are a component of AD pathology, perhaps related to β-amyloid biogenesis^39, 40^ or a cellular response to amyloid^41^. Interestingly, the extracellular droplets in human post-mortem β-amyloid plaques resembled the droplet-like architecture of ApoE- and premelanosomal protein-associated intraluminal vesicles, which are proteins necessary for the amyloid structure of retinal melanosomes^42^. Fragments of open lipid membrane were also observed that were absent from mouse models of FAD^29^ and control post-mortem tissue. Since neurodegeneration is absent in FAD mouse models, it is possible that the presence of membrane fragments is indicative of the cellular damage that culminates in neuronal loss in AD.

Aβ fibrils *in situ* were broadly consistent with earlier *ex vivo*, helically averaged atomic models, particularly type 1, 2, and type 1b^20^. However, additional amyloid species were also apparent, including protofilament-like rods and branched amyloid that were similar to those observed in cryo-sections and *ex vivo* purified amyloid from an FAD mouse model^29^. The existence of branched fibrils and protofilament-like rods within post-mortem AD brain is suggestive of catalytic mechanisms involving secondary nucleation-mediated fibril growth^43, 44^ and could offer an explanation for the focal concentration of Aβ observed in β-amyloid plaques^29^.

In-tissue tomographic volumes of tau pathology indicated filaments were unbranched and arranged in parallel clusters. These were observed within cells and in extracellular locations. The latter are consistent with tau clusters that remain after degeneration of the neurite^18^. Two distinct tau ultrastructural polymorphs, PHF and SF, are associated with AD^23, 45^. These differ in the symmetric versus asymmetric arrangements of protofilaments within an otherwise twisted filament of tau (2.5 and 2.2°.nm^−1^ twist of PHF and SF, respectively). Cryo-section and cryoFIB-SEM liftout lamella tomograms both produced subtomogram averaged maps that resolved the protofilament substructure of specific tau filament conformers *in situ,* that originated from distinct tissue locations. The subtomogram average structure of most tau clusters accommodated the atomic model of PHF^23^. However, these subtomogram averages exhibited structural variability of helical twist. The marked similarity of fibrils within the same cluster compared to those in different clusters suggest that the specific subcellular environment influences the structure of tau filaments. We did not obtain a subtomogram average that resembled SF in cryo-sections or liftout lamellae tomograms. This may reflect the lower proportion of SFs in AD brain^23, 24^.

Clusters of filaments resembling different tau conformers were also observed within tau threads at the periphery of an amyloid plaque. Subtomogram averaging of filaments within one of the tau threads revealed a map that best fit the atomic model of AD PHFs. The structure of a cluster of filaments within a second tau thread that neighboured the first did not match the atomic models of PHF or SF. Instead, the map most closely resembled the CTE type 1 filament^46^. We cannot rule out that this form of tau represents an additional conformer that has not yet been observed in *ex vivo* purified samples^47^, and must await higher resolution structure determination by cryoET to resolve this possibility. These data report the first incidence of a tau filament structure resembling the CTE conformer in AD brain. Multiple distinct tau filament conformers associated with different tauopathies converging in the same individual have been reported previously^28, 46^. However, here the post-mortem donor’s clinical record did not report chronic head trauma or impacts, and did not show co-pathology indicative of CTE or any neurodegenerative disease other than AD (**Extended Data Fig. 1**). Interestingly, CTE tau filaments have recently been identified in other neurodegenerative conditions aside from CTE, suggesting that this fold could be associated with neuroinflammation and a broad variety of environmental insults^47, 48^. Importantly, the distinct clusters of PHF and CTE-like tau filaments were spatially segregated, forming distinct amyloid deposits located ∼1 μm apart, within two separate microscopic regions of pathology enclosed by different subcellular compartments. This indicates that the local cellular environment is a major determinant of which tau filament structures are formed in the brain.

Application of the *in situ* structural workflows reported here to larger cohorts of diverse AD donors and across different brain regions, may further reveal how the spatial organisation of amyloid of different structures relates to individual neuropathological profiles. It will also be important to apply these approaches to other neurodegenerative diseases, many of which share related or overlapping types of amyloid neuropathology.

## SUPPLEMENTARY INFORMATION

**Extended Data Table 1.** Statistics and constituents of in-tissue tomograms from a MX04-labelled plaque and surrounding areas.

**Extended Data Table 2.** Statistics and constituents of in-tissue tomograms from MX04-labelled tau pathology and surrounding areas

**Extended Data Table 3.** Statistics and constituents of in-tissue tomograms from MX04-labelled control patient brain

**Extended Data Table 4.** Statistics and constituents of in-tissue cryo-liftout tomograms from MX04-labelled tau pathology and surrounding areas.

**Extended Data Table 5.** Parameters for cryoET data collection and subtomogram averaging of tau filaments

**Extended Data Table 6.** Parameters for cryoET data collection and subtomogram averaging of Aβ fibrils.

**Extended Data Figure 1.** Neuropathological characterisation of post-mortem AD donor.

**Extended Data Figure 2**. Biochemical and immunohistochemical profile, cryoFM targeting of MX04-labelled amyloid pathology of post-mortem AD donor tissue.

**Extended Data Figure 3.** Cryo-CLEM of MX04-labelled post-mortem AD brain.

**Extended Data Figure 4.** In-tissue cryoET of MX04-labelled β-amyloid plaque cryo-sections from fresh AD post-mortem brain donor.

**Extended Data Figure 5.** In-tissue cryoET of tissue cryo-sections from fresh non-demented control post-mortem brain donor.

**Extended Data Figure 6. I**n-tissue cryoET of MX04-labelled tau inclusion cryo-sections from fresh AD post-mortem brain donor.

**Extended Data Figure 7.** Measurement of fibril widths in raw tomographic maps, helical twist tau subtomogram averages and unsupervised classification of tau filament subvolumes.

**Extended Data Figure 8.** Subtomogram averaging of whole filaments (stalkInit see Methods) from tau clusters within tissue cryo-sections situated in six distinct locations, shown in panels a-e.

**Extended Data Figure 9.** Available atomic models of tau conformers fitted into the map and resolution estimation based on Fourier shell correlation (FSC) of half-maps from subtomogram averaging of tissue cryo-section tomograms.

**Extended Data Figure 10.** Available atomic models of known tau amyloid conformers fitted into the map and resolution estimation based on Fourier shell correlation (FSC) of half-maps from subtomogram averaging of different tau filament clusters a-f.

**Extended Data Figure 11.** Subtomogram averaging of Aβ fibrils from MX04-labelled β-amyloid plaque cryo-sections.

**Extended Data Figure 12.** Confocal cryoFM of MX04-labelled post-mortem AD brain.

**Extended Data Figure 13.** Correlated cryoFM-FIB-SEM liftout targeting of MX04-labelled tau pathology in high-pressure frozen post-mortem AD brain.

**Extended Data Figure 14.** CryoFIB-SEM liftout of MX04-labelled post-mortem AD brain containing tau

**Extended Data Figure 15.** Distinct structures of filaments from different tau clusters in cryoFIB-SEM liftout lamella tomograms.

**Extended Data Figure 16.** Subtomogram averaging scheme of tau pathology in tissue cryo-sections. Related to Fig. 3i-j.

**Extended Data Figure 17.** Subtomogram averaging scheme of tau pathology in tissue cryoFIB-SEM liftout lamella. Related to Fig. 4h-i.

**Extended Data Figure 18.** Subtomogram averaging scheme of MX04-labelled tau pathology in tissue cryoFIB-SEM liftout lamella. Related to Fig. 4j-k.

## METHODS

### Data reporting

No statistical methods were used to predetermine sample size. The experiments were not randomized and the investigators were not blinded to allocation during experiments and outcome assessment.

### Donor and ethical information

Post-mortem brain tissue was obtained through the Netherlands Brain Bank (NBB; Amsterdam, The Netherlands, https://www.brainbank.nl). In compliance with all ethical standards, brain donors signed informed consent regarding the usage of their brain tissue and clinical records for research purposes. The local medical ethics committee of the VUmc approved the brain donor programs of the NBB. Brain dissection and neuropathological diagnosis were performed according to international guidelines of Brain Net Europe II (BNE) consortium (http://www.brainnet-europe.org) and NIA-AA^49^.

Fresh, flash-frozen post-mortem AD and non-demented donor post-mortem brain were cryopreserved at –80°C and provided a source of tissue for these studies. The control case was a 90-year-old man with a history of depression and prostate cancer. At the age of 87, the donor was admitted to a nursing home. In the last phase, he was passive with a concentration disorder, but he was not demented, with normal language skills, speaking skills and communicative ability. Neuropathological examination revealed slight atrophy of the temporal lobe. No β-amyloid plaques or neurofibrillary tangles were observed in the temporal lobe.

### Neuropathology of donor tissue

A tissue block containing mid-temporal gyrus that was adjacent to the high-pressure frozen fresh tissue, was formalin-fixed and paraffin embedded. Sections of 5 µm thickness were prepared and mounted on Superfrost+ microscope slides (VWR, Amsterdam, The Netherlands). After overnight incubation at 37°C, slides were deparaffinized using xylene and alcohol and subsequently washed in phosphate buffered saline (PBS; pH 7.4).

For the histochemical detection of plaques and NFTS, the method described by Gallyas was used^50^. In short, tissue was pretreated using 5% w/v periodic acid for 30 min. Subsequently, the tissue was silver impregnated using a 0.035% w/v silver nitrate solution for 30 min. After silver impregnation, the bound silver was developed using a reduction reaction induced by the development solution (2.5% w/v sodium carbonate, 0.1% w/v silver nitrate, 0.5% w/v tungstosilicic acid hydrate, 0.1% w/v ammonium nitrate and 0.1% w/v formaldehyde). The development was stopped by washing in 0.5% w/v Acetic acid for 5 min and unbound silver was removed by washing in 5% w/v sodium thiosulfate for 5 min. Sections were counterstained using haematoxylin (Diapath, Martinengo, Italy). The sections were dehydrated using alcohol and xylene and coverslipped using Depex (BDH Laboratories Supplies, East Grinstead, UK).

For immunohistochemistry, deparaffinized sections were pretreated with 0.3% Hydrogen peroxide in PBS for 30 min to block endogenous peroxidase activity, followed by autoclave heating (121 °C for 20 min) in 10 mM sodium citrate buffer (pH 6) for antigen retrieval. Primary antibodies were incubated overnight at room temperature and diluted in antibody diluent (Sigma Aldrich, St. Louis, MO, USA) as follows: anti-pTauSer202/Thr205 clone AT8 (ThermoFisher Scientific, Waltham, MA, USA) 1:800, anti-amyloid beta clone 4G8 (Biolegend, San Diego CA, USI) 1:1000, pTau-Thr217 (ThermoFisher Scientific) 1:6400, P62-lck (BD Biosciences, CA) 1:1000, anti-alpha-synuclein (phospho S129) (Abcam, Cambridge, UK) 1:500, and anti-pTDP-43 Ser409/410 (Cosmo Bio USA, CA, USA) dilution 1:6000. Envision mouse/rabbit HRP (DAKO, Glostrup, Denmark) was used in the secondary detection step, and 3,3’-Diaminobenzine (DAB, DAKO) was used as a chromogen. Immunostained sections were counterstained using haematoxylin, dehydrated using alcohol and xylene, and coverslipped using Depex.

### Preparation of fresh post-mortem brain slices

100-200 μm slices of post mortem AD and control donor brain were sliced (speed 0.26 mm/s) using a vibratome (VT1200S, Leica) in ice-cold carboxygenated NMDG buffer (93 mM NMDG, 2.5 mM potassium chloride, 1.2 mM sodium hydrogen carbonate, 20 mM HEPES, 25 mM glucose, 5 mM sodium ascorbate, 2 mM thiourea, 3 mM sodium pyruvate, 10 mM magnesium sulphate heptahydrate, 0.5 mM calcium chloride dihydrate, pH 7.4, 300 - 315 mOsmol)^51^.

### Immunohistochemistry and confocal fluorescence microscopy

Free-floating (200 μm) acute brain slices were incubated in carboxygenated NMDG buffer to which 15 nM methoxy-X04 (MX04) was diluted for 1 h. Next, the slices were transferred to fresh NMDG buffer and fixed with 4% paraformaldehyde (PFA). Slices were permeabilised with 2% Triton X-100 for 30 mins, then incubated in blocking buffer (3% w/v BSA, 0.1% w/v TritonX-100, 50 mM Tris-Cl, 150 mM NaCl, pH7.4) for 1 h at room temperature. After every incubation the tissue sample was washed 3x with TBS (50 mM Tris-Cl, 150 mM NaCl, pH7.4). To detect β-amyloid and tau inclusions, slices were incubated in 1:750 dilution 6E10 (Biolegend, 803001) or 1:750 dilution 4G8 (Biolegend, 803001) and 1:750 dilution AT8 (Thermofisher) in blocking buffer at 4°C for 24 h, respectively. After 3 x washes with TBS for 5 min, slices were incubated in 1:1000 diluted anti-mouse IgG1 AF-633 (Thermo Fisher, A21126) or 1:1000 diluted anti-mouse IgG2BAF-633 (Thermo Fisher, A21126) and 1:1000 diluted anti-mouse IgG1 AF-568 (Thermo Fisher) in blocking buffer for 2 h at room temperature. Following 3 washes in TBS for 5 min each, slices were mounted with Vectashield (Vector Laboratories) on a microscope slide (Erpredia, J1810AMNZ). Images were acquired with a confocal laser scanning microscope (ZEISS LSM 700) using a 10x/0.3 and a 63x/1.4 numerical aperture (NA) air objective lens, with frame size 1024×1024 pixels and 512×512 pixels, respectively. MX04, AF-568, and AF-633 were detected excitation/emission maxima of 405/435 nm, 579/603 nm, and 639/669 nm, respectively.

### Sarkosyl-insoluble tau purification and immunoblots

Sarkosyl-insoluble tau purification followed a previously published protocol^52^. Briefly, fresh post-mortem brain tissue was homogenized in 10 vol (w/v) of homogenisation buffer (10 mM Tris-HCI (pH 7.4), 0.8 M NaCI, 1 mM ECTA, 10% w/v sucrose). The homogenate was centrifuged at 20,000 x *g* for 20 min, at 4°C and the supernatant was retained. The pellet was re-homogenized in 5 vol (w/v) of homogenization buffer and re-centrifuged. Both supernatants were combined, brought to 1% N-lauroylsarcosinate (w/v), and were centrifuged at 100,000 x g for 1 h at 21°C. The sarkosyl-insoluble pellets were resuspended in 50 mM Tris, pH 7.4 (0.2 ml per g of starting material) and stored at 4°C for immunoblots. Samples were run using 4–12% Bis-Tris-Gel (Thermo fisher) and transferred onto PVDF membrane using iBlot gel transfer stacks (Thermo Fisher). The PVDF was blocked (2.5 % w/v casein in 0.1% Tween 20, 50 mM Tris.Cl pH7.4, 100 mM NaCl) for 1 h at room temperature. The following primary antibodies were diluted in 1.25% w/v casein in TBS-T (0.1% Tween 20, 50 mM Tris.Cl pH7.4, 100 mM NaCl): 1:2000 Tau 46 (aa 404-441, T9450, Merck); 1:1000 AT8 (pS202/pT205 Tau, MN1020, Thermo Fisher); 1:1000 4-repeat tau (aa 275-291, 05-804, Merck); 1:500 3-repeat tau (aa 267-316, 05-803, Merck). The PVDF membranes were incubated with primary antibodies at 4°C overnight. The membranes were washed five times with TBS-T for 5 min, followed by incubation of secondary antibody for 40 min at room temperature, then washed five times in TBS-T for 5 min. PVDF membranes with ECL reagent (Lumigen) were imaged on an iBright 1500 (Thermo Fisher).

### High pressure freezing

Acute brain slices were incubated in carboxygenated NMDG buffer, to which 15 nM methoxy-X04 (MX04) was diluted for 1 h at room temperature. Next, the slices were washed three-times in carboxygenated NMDG for 5 min each. Grey matter biopsies with a diameter of 2 mm were incubated in cryoprotectant (5% w/v sucrose and 20% w/v 40,000 Dextran in NMDG buffer) for 30 min at room temperature. 100 μm deep wells of the specimen carrier type A (Leica, 16770152) were filled with cryoprotectant, and the tissue biopsies were carefully placed inside to avoid tissue damage. They were then covered with the flat side of the lipid-coated specimen carrier type B (Leica, 16770153) and high-pressure frozen (∼2000 bar at −188°C) using a Leica EM ICE.

### Cryogenic fluorescence microscopy (cryoFM)

High pressure frozen samples were imaged using a cryogenic fluorescence microscope, Leica EM Thunder with a HC PL APO 50x/0.9 NA cryo-objective, Orca Flash 4.0 V2 sCMOS camera (Hamamatsu Photonics) and a Solar Light Engine (Lumencor) at −180°C. A DAPI filter set (excitation 365/50, dichroic 400, emission 460/50) was used to detect MX04-labelled amyloid. A rhodamine filter set (excitation 546/10, dichroic 560, emission 525/50) was used as a control imaging channel. The images were acquired with a frame size of 2048×2048 pixels. Tile scans of high-pressure frozen carriers were acquired with 17% laser intensity for 0.1 s. *z*-stacks of ultrathin cryo-sections were acquired with 30% intensity and an exposure time of 0.2 s. Images were processed using Fiji ImageJ.

### Cryo-ultramicrotomy

High pressure frozen sample carriers were transferred to a cryo-ultramicrotome (Leica EM FC7, −160°C) equipped with trimming (Trim 20, T399) and CEMOVIS (Diatome, cryo immuno, MT12859) diamond knives. A trapezoid stub of tissue measuring 100 x 100 x 60 μm was trimmed, which contained the target amyloid. Cryo-sections (70 nm thick) were then cut at −160°C with a diamond knife (Diatome, cryo immuno, MT12859) and adhered onto a glow discharged (Cressington glow discharger, 60 s, 10-4 mbar, 15 mA) 3.5/1, 300 mesh Cu grid (Quantifoil Micro Tools) using a Crion electrostatic gun and gold eyelash micromanipulators.

### Cryogenic correlated light and electron microscopy (cryo-CLEM) of cryo-sections

The location of amyloid plaques in tissue cryo-sections was assessed by cryogenic fluorescence microscopy based on Methoxy-X04 fluorescence (excitation 370 nm, emission 460-500 nm).. Grid squares that showed a signal for MX04 were selected for cryoET. The alignment between cryoFM images and electron micrographs were carried out using a Matlab script^53, 54^, in which the centres of 10 holes in the carbon foil surrounding the region of interest were used as fiducial markers to align the cryoFM and cryoEM images.

### cryoCLEM and focussed ion beam scanning electron microscopy (cryoFIB-SEM) of liftout lamellae

Carriers containing high pressure frozen samples for cryoFIB-SEM liftout were transferred to a cryo-ultramicrotome (Leica EM FC7, −160°C) equipped with a Trim 45 T1865 diamond knife. The surface of the carrier and high-pressure frozen tissue within were trimmed to remove surface ice contamination and the top layer of vibratome-damaged tissue^55^. To achieve this, three 300 μm wide steps were trimmed back on 4 sides of the carrier: the outermost step was trimmed ∼30 μm deep, the next was trimmed ∼20 μm back, and the innermost step was trimmed ∼10 μm back. The resulting protruding square of tissue was then trimmed ∼2-5 μm back to achieve a smoother less contaminated surface for cryoFIB-SEM and liftout.

The cryoCLEM workflow was performed on a ZEISS Crossbeam 550 Focused Ion Beam-Scanning Electron Microscope (FIB-SEM) equipped with the Quorum cryo-system, ZEISS Cryo-accessories tool kit and Omniprobe 350 cryo-micromanipulator (Oxford Instruments) and operated at 30 keV. For cryogenic fluorescence microscopy (cryoFM) a ZEISS LSM (laser scanning microscope) 900 based on an upright Axio Imager stand equipped with Airy Scan and Linkam cryo-stage was used.

At the Quorum prep desk, the high pressure freezing (HPF) sample carrier (Leica Microsystems, Vienna, Austria) was mounted on the corresponding ZEISS universal sample holder (USH, ZEISS cryo accessories tool kit). First, the ZEISS USH was placed on the ZEISS adapter for the Linkam cryo stage used for performing cryoFM. The assembly was transferred into the Linkam cryo stage in liquid nitrogen using the ZEISS transfer box. A LM ZEN Connect project was acquired using the plug-in ZEN Connect of the ZEN Blue software (Zeiss Microscopy, Jena, Germany). One overview image of the sample and the holder was acquired with a 5x, 0.2 NA C-Epiplan Apochromat air objective with an Axiocam 503. MX04, reflection (for correlation with the EM images) and a control channel were imaged with a 385/511/567 nm LED, respectively, in combination with a quad-band filter (Excitation BP 385/30, BP 469/38, BP 555/30, BP 631/33, Emission QBP 425/30 + 514/30 + 592/25 + 709/100). The image was acquired with a frame size of 2.79 x 2.10 mm and 1936×1460 pixels, 5/3/60% LED power and 50/6/300 ms exposure time for MX04/reflection/control.

For sample quality assessment a large 3D volume (1.27 x 1.27 x 0.108 mm/ 2824×2824 pixels, 49 z-slices) was scanned with a 10x, 0.4 NA C-Epiplan Apochromat air objective in confocal mode. MX04 (0.04 % laser power, 405 nm laser, detection window 410-546 nm, pixel dwell time 0.74 µs) and the reflection (0.01% laser power, 640 nm laser, detection window 630-700 nm, pixel dwell time 0.74 µs) were imaged.

Regions of interest were scanned with a 100x, 0.75 NA LD EC Epiplan-Neofluar air objective (125.15 x 125.15 x 19.8 µm/ 1140 x 1140 pixels, 37 z-slices, pixel dwell time 1.8 µs) with the following settings: MX04 (0.2 % laser power, 405 nm laser, detection window 410-544 nm), reflection (0.05% laser power, 640 nm, detection window 639-700 nm). A linear deconvolution was run on all confocal images (10x and 100x) using the ZEISS LSM plus plugin.

Airyscan SR (super-resolution) *z*-stacks were acquired from the regions of interest using the Airyscan 2 detector and the 100x objective detailed above. MX04 was detected using the 405 nm laser line (1.5 % laser power, 405 nm laser, pixel dwell time 2.18 µs, detection window 400-650 nm, image size: 62.31 x 62.31 x 13.65 µm/ 946 x 946 pixels, 36 z-slices). Airyscan images were processed using the Airyscan joint deconvolution (jDCV) plugin of the ZEN blue software. CryoCLEM alignment was performed with maximum projections of confocal and Airyscan image *z*-stacks prepared in ZEISS ZEN blue (version 3.6). After finalising light microscopy, the USH connected to the Linkam adapter was transferred into the Quorum Prep box. The USH was detached from the Linkam adapter and mounted on the ZEISS cryo lift-out sample holder with the USH in flat orientation and a mounted upright, standing half-moon Omniprobe grid clipped into an AutoGrid (Thermo Fisher). Using the Quorum cryo-shuttle the sample holder was transferred into the Quorum prep chamber attached to the Crossbeam 550 FIB-SEM. The temperature of the cryo-stage/anti-contaminator in the main and prep chamber was set to −160°C/−180°C, respectively. The sample was sputter-coated with platinum for 45 seconds (5 mA current). After the sputter-coating the sample was transferred on the Quorum cryo stage in the main chamber.

CryoFM-targeted cryoFIB-SEM liftout was carried out driving the ZEISS Cross beam FIB-SEM stage within a ZEN Connect imaging project, in which cryoFM, SEM and FIB images were correlated using surface features (cryoplaning markings and ice contamination) as fiducial markers (**Extended Data Figs. 11, 12 & 13**)^56^. At normal view (0° stage tilt), 5 mm working distance and 2.3 kV acceleration voltage an overview of the HPF carrier and a zoomed-in SEM image were acquired and loaded into a new SEM session of the existing ZEN Connect project (imaging parameter: SE2(Everhart-Thornley) detector, 98 pA SEM current, 4096 x 3072 pixels, 800 ns dwell time, line average with 23 iterations and 800/420 nm pixel size, respectively. The SEM session was correlated with the already acquired cryoFM session by using the reflection mode channel images for alignment.

Since image navigation was desired in FIB mode, the stage was tilted to 54° allowing normal FIB view. The sample was brought into the coincidence point of SEM and FIB with a 5 mm working distance and an overview FIB image (imaging current 50 pA, SE2 detector, 2048 x 1536, 1,6 µs dwell time, pixel average, 300 nm pixel size) was taken for FIB session alignment with the former SEM/cryoFM session. The coincidence point was fine adjusted to the region targeted for liftout and the milling box for coarse cross-sectioning was positioned based on the alignment between cryoFM and FIB image. A ∼80 μm wide, 35 µm high and 30 μm deep trapezoidal cross-section was milled from the front side using a 30 nA FIB probe. Since ice contamination, especially on top of region of interest was observed, the sample holder was transferred back into the Quorum prep box for cleaning. Under cryogenic conditions the sample surface was cleansed by using a brush and by wiping using a swab. After an additional sputter-coating in the Quorum prep chamber (see above for parameters), a cold deposition of platinum precursor was applied in the Crossbeam main chamber. For cold deposition, the distance between sample and gas injection capillary was about 3 mm and the gas reservoir valve was opened for 45 sec. The gas reservoir temperature was about 28 °C (unheated gas reservoir state). Using the saved stage position the cross-section at the region of interest was targeted, the coincidence point alignment was checked. A FIB image was taken and aligned with the former FIB session using the already milled cross-section as reference.

Next, a 30 nA FIB probe was used to mill a second corresponding cross-section from the back side and a ∼60 μm wide left side cut that left a ∼80 μm wide, 10 μm thick, 30 μm deep tissue chunk attached on to its right side. At the front side the cross-section was additionally polished using a 15 nA FIB probe. The stage was tilted to 10° tilt for milling a ∼80 μm wide L-shaped undercut, leaving a small connection on the left side (7nA FIB probe). As a liftout tool, a ∼5 µm thick copper block was attached to the Omniprobe manipulator tip. The stage was at 10° tilt to allow access of the micromanipulator whilst bypassing the Autogrid ring. Before liftout the ∼5 μm copper block was attached to the right side of the tissue chunk using redeposition milling of copper material (three 2.5 x 5 μm milling windows with 700 pA FIB probe and 140 mC/cm² dose). Next, the tissue chunk was cut free from the left side to achieve liftout. At 10° stage tilt the ∼80 μm wide chunk was attached by redeposition milling to a half-moon EM grid (Omniprobe) clipped into an AutoGrid (ThermoFisher) and cut in half, leaving the distal ∼40 μm chunk attached to the EM grid. The remaining proximal ∼40 μm chunk attached to the Omniprobe, was attached to a second location on the EM grid. The stage was tilted to 56° before two 8-10 μm wide, ∼15 μm deep lamella windows were milled in each chunk half. At 56° stage tilt, the angle between tissue chunk/grid plane and FIB beam is 2° allowing to bypass the outer AutoGrid ring for lamellae thinning. The lamellae were sequentially thinned from both sides to ∼2 µm, then ∼1 um, then ∼500 nm and finally 130 to 200 nm using 700 pA, 300 pA, 100 pA and 50 pA FIB probes, respectively. Each lamella window was framed with unmilled tissue at the left, right and bottom side.

### Cryo-electron tomography and image reconstruction

Cryo-electron tomography was performed using a ThermoFisher 300 keV Titan Krios G2, X-FEG equipped with a Falcon4i detector and Selectris energy filter in the Astbury Biostructure Laboratory at the University of Leeds. A dose symmetric tilt scheme^57^ was implemented using Tomo5.12 software (ThermoFisher) to collect tilt series from −60° to +60° in 2° increments with a 100 μm objective aperture and 5-6.5 μm defocus. Each tilt increment received ∼2.3 s of exposure (fractionated into 8 frames) at ∼2 e-/Å^2^ per tilt for a total dose of ∼120 e-/Å^2^ per tilt series with a pixel size of 2.4 Å. Frames were aligned using motionCor2^58^ and tomograms were reconstructed from aligned tilt series using AreTomo^59^. For segmentations and preparing tomographic slice figures tomograms were deconvolved with Isonet^60^.

### Subtomogram averaging

Subtomogram alignment and averaging of Aβ fibrils and tau filaments was initially performed on a per-tomogram basis to assess the relationship between amyloid structure and its subcellular context. Because of the dense architecture of in-tissue amyloid, only tomograms with axial (oriented in the *z*-axis of the reconstructed tomogram) fibrils and filaments provided sufficient contrast to accurately pick individual fibrils and filaments for subtomogram averaging. Tilt series used for subtomogram averaging of Aβ fibrils or tau filaments were aligned in IMOD (v4.12.35)^61, 62^ using patch tracking, CTF-corrected and reconstructed by weighted-back projection. See **Extended Data Table 5** and **6** for per-tomogram subtomgram averaging details of tau filaments and Aβ fibrils, respectively.

For tau filaments, subtomogram averaging was performed on 9 cryoET volumes (7 from cryo-sections and 2 from liftout lamellae). Between 50 and 257 tau filaments (24×100×24 voxels) were manually picked from 4x binned (9.6 Å voxel size) tomographic reconstructions in IMOD. Coordinates of each filament were represented as a two-point contour with respective ‘head’ and ‘tail’ model points positioned at the poles of each filament (**Extended Data Figs 16a-b, 17a-b and 18a-b**). The slicer function of IMOD was used to rotate and translate each tau filament, ensuring different models were approximately centered along the filament axis. Using these model files as input, a script invoking the PEET ‘stalkInit’ command was run in default mode to generate new, single-point model files (containing coordinates of the head, centroid and tail), initial motive lists (MOTL) and rotation axes files, to supply for alignment and averaging in particle estimation for electron tomography (PEET1.16)^63, 64^. To minimize alignment to the missing wedge and to verify the accuracy of model point coordinates, initial averages were generated by restricting rotational and translation searching from the centroid of the *de novo* reference filament axis (**Extended Data Figs 16c, 17c and 18c**). In subsequent PEET alignment iterations, rotational and translational searching was performed, using the updated centroid coordinates and orientations as input. Where visible improvements in map resolution were observed, createAlignedModel was run to generate new models, motive lists and rotation axes containing the updated locations and orientations of particles from the alignment. Cylindrical masks with blurred edges (**Extended Data Figs 16d, 17d and 18d**) and then custom masks (generated in IMOD using imodmop) were applied to the stalkInit reference volume during subvolume alignment (**Extended Data Figs 16e, 17e and 18e**).

To improve alignment further, centroid models were added every 40 voxels along the stalkInit filament axis (9509 new model coordinates from the 9 cryoET volumes) using the ‘addModPts’ command. AddModPts models were used as new inputs for subvolume alignment and averaging in PEET (**Extended Data Figs 16f, 17f and 18f**). CreateAlignedModel was also run after addModPts alignments that generated improved map resolution, outputting new models and MOTLs of the updated locations and orientations of particles. After alignment iterations with 4x binned tomographic reconstructions, the ‘imodtrans’ command was used to generate scaled model files (96×40×96 or 104×40×104 voxel box size) for aligning with unbinned tomographic reconstructions (2.4 Å voxel size). Cylindrical masks with blurred edges (**Extended Data Figs 16g, 17g and 18g**) and then custom masks were applied to the addModPts reference volumes during alignment (**Extended Data Figs 16h, 17h and 18h-i**).

For both stalkInit and addModPts averages, the references used for alignment and averaging of subvolumes were progressively filtered to higher spatial frequency cutoffs in final iterations of alignment. To avoid overfitting, the spatial frequency cutoff applied to the reference was verified by Fourier Shell Correlation (FSC), to ensure the reference was not filtered to resolutions exceeding the FSC 0.5 criterion. Final stalkInit and addModPts subtomogram averages from each tomogram were generated from 40-121 and 494-2534 model points, respectively (**Extended Data Table 5**).

For Aβ fibrils, subtomogram averaging was performed on the two cryo-section volumes collected from MX04-labelled β-amyloid plaques that contained axially oriented fibrils. The same manual picking and subtomogram averaging procedures were followed (using stalkInit) as described for tau filaments. Initially, 100 fibrils were picked from one tomogram, with fibrils represented as two-point ‘head’ and ‘tail’ models, positioned at opposite ends of each fibril. This model was used as input to the PEET ‘stalkinit’ command to generate new model files (head, centroid, tail), initial motive lists, and rotation axes files, for alignment and averaging in PEET. The subtomogram average of 100 fibrils generated a featureless, smooth tube, with reducing detail observed with increasing fibril numbers used for averaging, a possible indication of particle heterogeneity (**Extended Data Fig. 11a**). To assess fibril heterogeneity, the widths of individual fibrils were manually measured in IMOD. On the basis of average fibril width, three subpopulations of fibrils were picked (3-5 nm protofilament-like rods, 4-9 nm fibrils and 6-12 nm wide fibrils). Subtomogram averaging of 20 protofilament-like rods, 42 fibrils, and 42 wide fibrils from two tomograms was performed with PEET as before (**Extended Data Fig. 11b-d** and **Extended Data Table 6**).

To compare the structural similarity of *in situ* subtomogram averages with *ex vivo* purified tau conformers, available atomic models were fitted in ChimeraX^65, 66^ (see **Extended Data Figs. 8, 9, 10** and **15**). Segmenting membranes within tomograms was performed in Dynamo^67^, using the manual surface modelling tool. Dynamo tables containing coordinate information of the tomogram membrane models were converted into CMM files and visualized in ChimeraX. The neural network-based tomogram segmentation pipeline in EMAN2^68^ was used to segment Tau filaments and Aβ fibrils.

### Annotation of constituents in tomographic volumes

The constituents of tomographic volumes of tissue cryo-sections from β-amyloid plaques, tau pathology, non-demented control, and cryoFIB-SEM liftout of tau threads were detailed in **Extended Data Tables 1, 2, 3** and **4**, respectively. Constituents were initially identified by two curators independently. Next, two curators consolidated and verified each annotation. The boundary of intracellular and extracellular regions of tomographic volumes were determined by the presence of myelinated axons or by lipid bilayers enclosing intracellular organelles within each tomographic volume and the corresponding electron micrograph used for cryoCLEM. The following constituents were identified: (1) Amyloid fibrils or filaments were assigned based on methoxy-X04 cryoCLEM labelling and rod shape. Fibrils and filaments were cross-checked for their absence in tomographic volumes reconstructed from non-demented control brain donors. (2) Vesicles were defined as closed spheroidal membranes. Extracellular vesicles were defined as membranes that were closed and situated within extracellular locations. (3) C-shaped vesicles: defined as cup-shaped membrane within the lumen of a vesicle. (4) Lipid droplets: 30–250 nm amorphous and smooth cuboidal or spheroidal particles^69^. (5) Extracellular droplets: amorphous and smooth cuboidal particles that were situated in extracellular locations. (6) Multilamellar bodies: defined as 40–250 nm vesicle or subcellular compartment wrapped in 3 or more concentric rings of 4.5-6 nm membrane lipid bilayer. (6) Mitochondria: defined by the double membrane including outer membrane and inner mitochondrial cristae. (7) putative F1Fo-ATPases: were identifiable within the inner mitochondrial membrane (for example see **Fig. 2c**). (8) Subcellular compartments: defined by plasma membrane containing membrane-bound organelles or a higher tomographic density than the extracellular space (excluding amyloid fibrils) that is consistent with the higher concentration of proteins in the cell cytoplasm compared to the extracellular space. (9) Actin: defined as ∼7 nm diameter filaments composed of a helical arrangement of globular subunits^70^. Actin was only observed intracellularly in a minor subset of both AD and non-demented control donor tomographic volumes. (10) Open membrane sheets: defined by ∼5 nm thick membrane lipid bilayers that did not form closed compartments. (11) Myelinated axon: defined as five or more layers of 6-8 nm membrane lipid bilayer enclosing a subcellular compartment^71^. (10) Knife damage: tissue cryo-sections contained small regions in which the sample has been compressed, leaving a crevasse in the tissue that were readily identified as holes within the tissue^34^.

### Data availability

Upon publication the subtomogram average data (refined maps, half-maps) will be deposited at the Electron Microscopy Data Bank (EMDB) under accession codes EMD-XXXX [https://www.ebi.ac.uk/pdbe/entry/emdb/EMD-XXXX] (AD Aβ protofibril), EMD-XXXX [https://www.ebi.ac.uk/pdbe/entry/emdb/EMD-XXXX] (AD Aβ type 1 fibril), EMD-XXXX [https://www.ebi.ac.uk/pdbe/entry/emdb/EMD-XXXX] (AD Aβ type 1b fibril), EMD-XXXX [https://www.ebi.ac.uk/pdbe/entry/emdb/EMD-XXXX] (AD Tau PHF), and EMD-XXXX [https://www.ebi.ac.uk/pdbe/entry/emdb/EMD-XXXX] (AD Tau CTE-like).

Upon publication all raw tomographic and cryoCLEM datasets will be deposited at Electron Microscopy Public Image Archive (EMPIAR) under accession codes EMPIAR-XXXXX [https://www.ebi.ac.uk/empiar/EMPIAR-XXXXX] (AD b-amyloid plaque cryo-sections), EMPIAR-XXXX [https://www.ebi.ac.uk/empiar/EMPIAR-XXXXX] (AD tau tangle cryo-sections), EMPIAR-XXXXX [https://www.ebi.ac.uk/empiar/EMPIAR-XXXXX] (non-demented control cryo-sections), and EMPIAR-XXXXX [https://www.ebi.ac.uk/empiar/EMPIAR-XXXX] (AD tau tangle cryolift lamellae).

## ACKNOWLEDGEMENTS

R.A.W.F. acknowledges UKRI Future Leader Fellowship (MR/V022644/1) and a University of Leeds Academic Fellowship. SER holds a Royal Society Professorial Fellowship (RSRP\R1\211057). The Astbury Biostructure Laboratory Titan Krios microscopes were funded by the University of Leeds and Wellcome Trust (108466/Z/15/Z & 221524/Z/20/Z). The Leica EM ICE, UC7 ultra/cryo-ultramicrotome and cryoCLEM systems were funded by Wellcome Trust (208395/Z/17/Z). We are grateful to the donors of post-mortem tissue, clinical records and the Netherlands Brain Bank. We thank Neil Ranson, Martin Wilkinson, and Anthony Turner (University of Leeds) for helpful discussion of amyloid structural biology. We thank Rebecca Thompson, Emma Hesketh, Louie Aspinall, Joshua White, and Oksana Degtjarik for help maintaining and setting up the Astbury Biostructure Laboratory (ABSL) cryoEM facility, including high-pressure freezing, cryoCLEM and Titan Krios microscopes. We thank Ruth Hughes and Sally Boxall (University of Leeds Bioimaging Facility) for support with fluorescence imaging experiments.

## Author Contributions

JJMH and THJM arranged donor tissues and performed neurohistopathological characterisation of post-mortem tissue. NF performed immunofluorescence and biochemical characterisation of post-mortem tissue and sample preparation. TJO, MAGG, NF, and MGe carried out cryoFM. TJO, MAGG, NF and RAWF carried out cryo-ultramicrotomy. AS performed cryoFIB-SEM liftout. MAGG and YH collected cryoEM and cryoET data. MAGG, TJO, JJ, and RAWF reconstructed cryoET data. MAGG and NF performed cryoCLEM analysis. NF, MAGG and RAWF annotated constituents of tomographic data. JJ, MAGG and NF carried out segmentations. JJ and MAGG performed subtomogram averaging. RAWF and SER supervised MAGG. RAWF devised and supervised the entire project. All authors contributed to writing the manuscript.

## Competing interests

The authors have no competing interests as defined by Nature Portfolio, or other interests that might be perceived to influence the results and/or discussion reported in this paper.

## Notes

### Competing Interest Statement

The authors have declared no competing interest.

