## Extended Data Figures 1-18 for "*In situ* cryo-electron tomography of β-amyloid and tau in post-mortem Alzheimer’s disease brain"

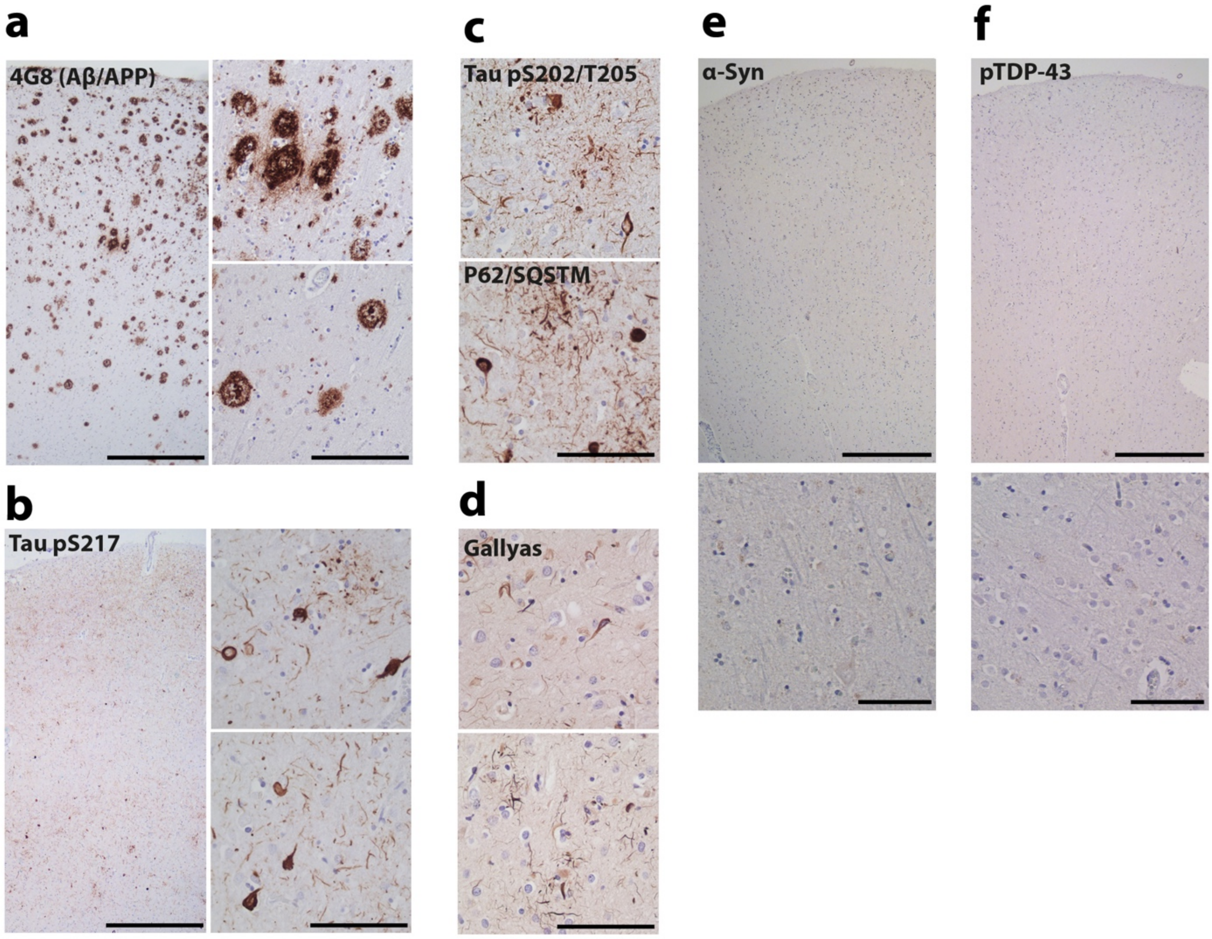


**Extended Data Figure 1. Neuropathological characterisation of post-mortem AD donor.** Formalin-fixed paraffin embedded tissue from the mid-temporal gyrus of the AD case was assessed for β-amyloid and phospho-Tau pathology by immunohistochemistry and silver staining. Cell nuclei were counterstained with hematoxylin.

**a** Left, overview showing β-amyloid deposits detected with 4G8, which were present throughout the cortical area from the pial surface to the subcortical white matter. Scale bar, 300 µm. Top right, close-up showing diffuse and compact plaques, as well as coarse-grained plaques. Bottom right, close-up showing dense-cored plaques. Scale bar, 100 µm.

**b** The presence of tau pathology was assessed with Tau phospho-S217 antibody, showing neurofibrillary tangles (NFT), neuropil threads (NT) and neuritic plaques (NP) that were present throughout the cortical region from the pial surface to the white matter. Left and right, overview and close-ups, respectively. Scale bars, 300 and 50 µm, respectively.

**c** Top and bottom, detection of Tau phosphoS202/T205 (AT8) and P62/SQSTM1 (a marker of aggregated protein) also revealed NFTs, NTs and NPs, respectively. Scale bar, 50 µm.

**d** Top and bottom, close-ups confirming the presence of amyloid in NFTs, NTs and NPs with Gallyas silver staining. Scale bar, 50 µm.

**e** and **f** α-synuclein and phospho-TDP-34 inclusions were absent. Top and bottom, overview and close-up, respectively. Scale bars, 300 and 50 µm, respectively.

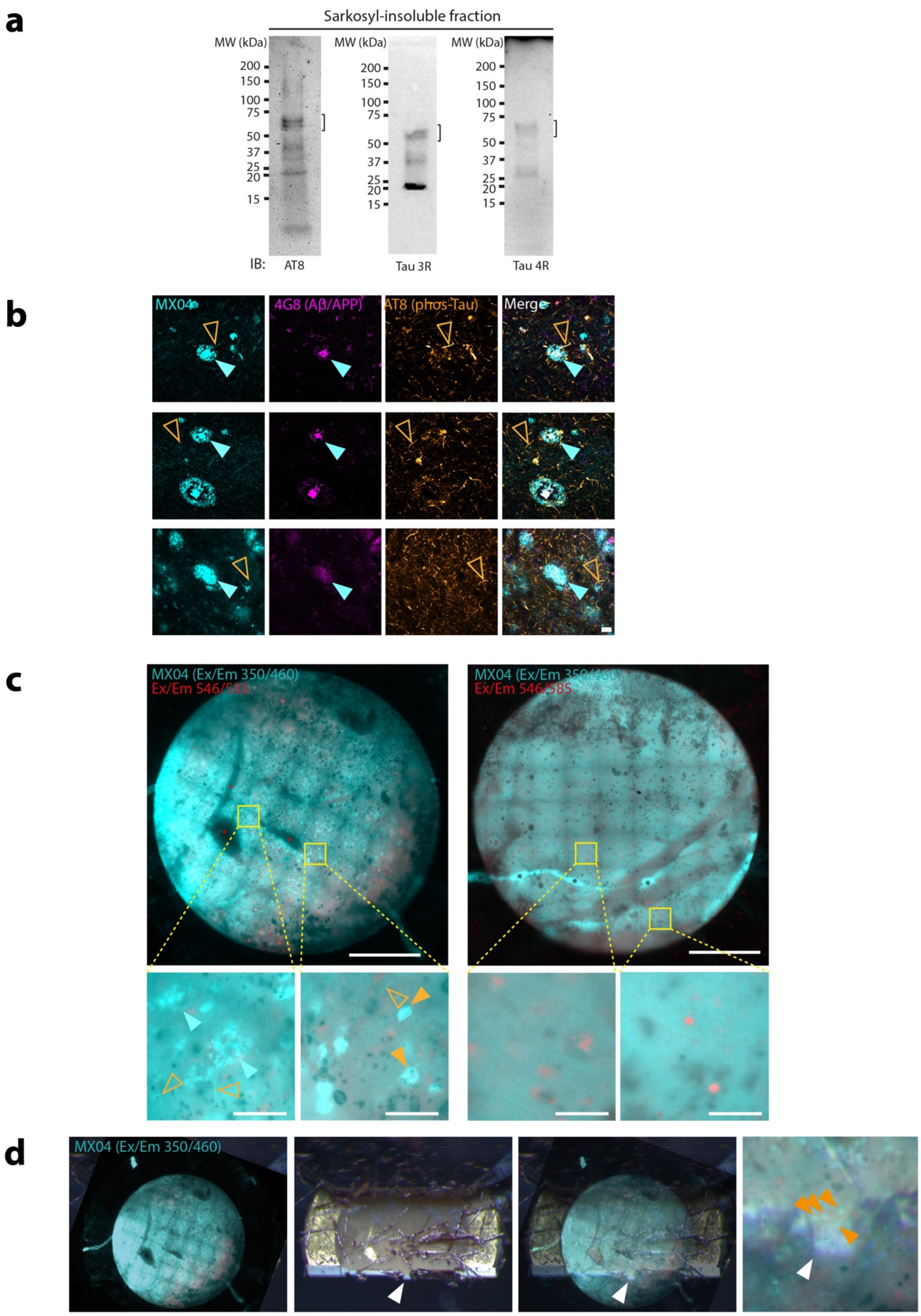


**Extended Data Figure 2. Biochemical and immunohistochemical profile, cryoFM targeting of MX04-labelled amyloid pathology of post-mortem AD donor tissue.**

**a** Immunoblot detection of sarkosyl-insoluble fraction of AD post-mortem brain. Left panel, probing with phospho-Tau (AT8, **see Methods**). Middle panel, 3-repeat (3R) tau immunoblot. Right panel, four-repeat (4R) tau immunoblot. Brackets, full-length tau bands.

**b** Related to **Fig. 1c**. Fluorescence confocal microscopic detection from left to right, amyloid (MX04), Aβ (4G8), phospho-Tau (AT8), and merged in AD post-mortem donor brain. Right, merged. Cyan arrowhead, β-amyloid plaque. Cyan arrowhead, β-amyloid plaque. Open orange arrowhead, tau thread. Top, middle and bottom, representative regions of Mid-temporal gyrus. Scale bar, 20 μm.

**c** CryoFM image of high-pressure frozen fresh post-mortem brain biopsy from AD (left) and non-demented control (right) donors. Cyan, MX04 fluorescence. Red, autofluorescence detected with excitation and emission of 546 nm and 585 nm, respectively. Yellow rectangle, regions shown as close-ups. Scale bar, 0.5 mm. Lower left panel, close-up showing putative β-amyloid pathology (cyan arrowhead) and tau threads (open orange arrowhead). Lower second from left panel, close-up showing putative tau tangle (closed orange arrowhead) and thread (open orange arrowhead). Scale bar, 50 μm.

**d** CryoFM targeting of cryo-ultramicrotomy. Left panel, cryoFM image of planchette containing MX04-labelled high-pressure-frozen tissue. Middle left panel, stereomicroscope image of the planchette during trimming with a cryo-ultramicrotome. White arrowhead indicates trapezoid stub of tissue targeted for the collection of cryo-sections. Middle right panel, alignment of cryoFM and stereomicroscope images. Right panel, close-up image of trimmed planchette with cryoFM image. Orange arrowhead, MX04-labelled amyloid within tissue stub (related to **Fig. 1f**).

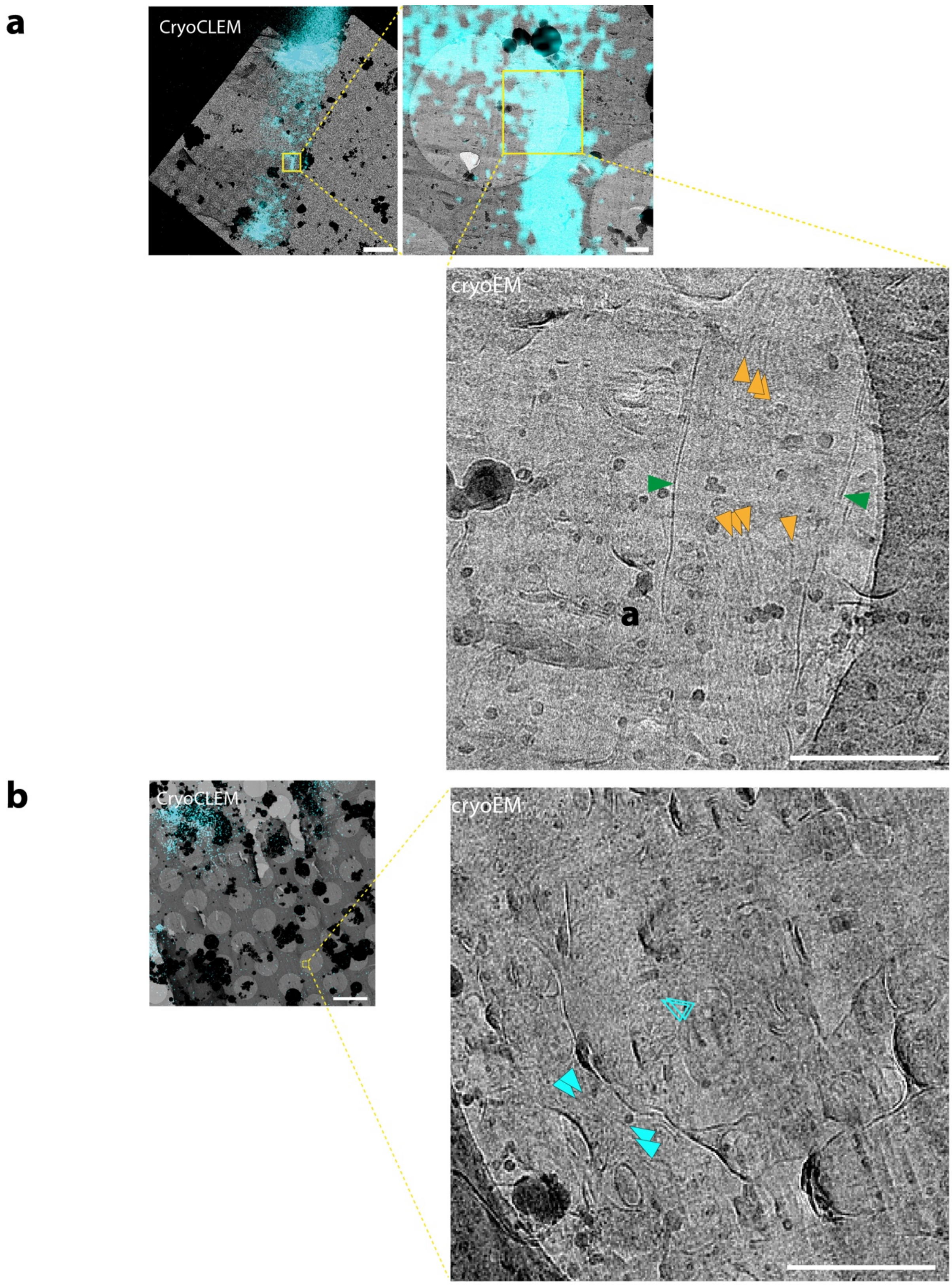


**Extended Data Figure 3. Cryo-CLEM of MX04-labelled post-mortem AD brain.**
**a** CryoCLEM of MX04-labelled tau inclusion within AD post-mortem brain cryo-section. Upper left panel, aligned cryoFM image (cyan, MX04) with cryoEM image. Yellow rectangle, area shown in close-up. Scale bar, 5 µm. Upper right panel, close-up. Yellow rectangle showing location of MX04-labelled neurite. Scale bar, 500 nm. Lower panel, medium magnification

cryoEM image of neurite containing MX04. Orange arrowheads, putative tau filament. Green arrowhead, putative plasma membrane of neurite. Scale bar, 500 nm. See cryoET data in **Extended Data Figure 6**)

**b** CryoCLEM of AD post-mortem brain cryo-section showing unlabelled amyloid deep in the tissue below the depth of MX04 penetration. Left panel, aligned cryoFM image MX04 with cryoEM image showing MX04 only labels top ~15 μm of 100 μm thick tissue biopsy. Yellow rectangle, region in cryo-section corresponding to 27 μm deep within the tissue biopsy that lacked MX04-label. Scale bar, 5 μm. Right panel, close-up of medium magnification cryoEM image showing region from which cryoET data were collected (see **Extended Data Figure 4c**). Cyan arrowheads, putative Aβ fibrils. Scale bar, 500 nm.

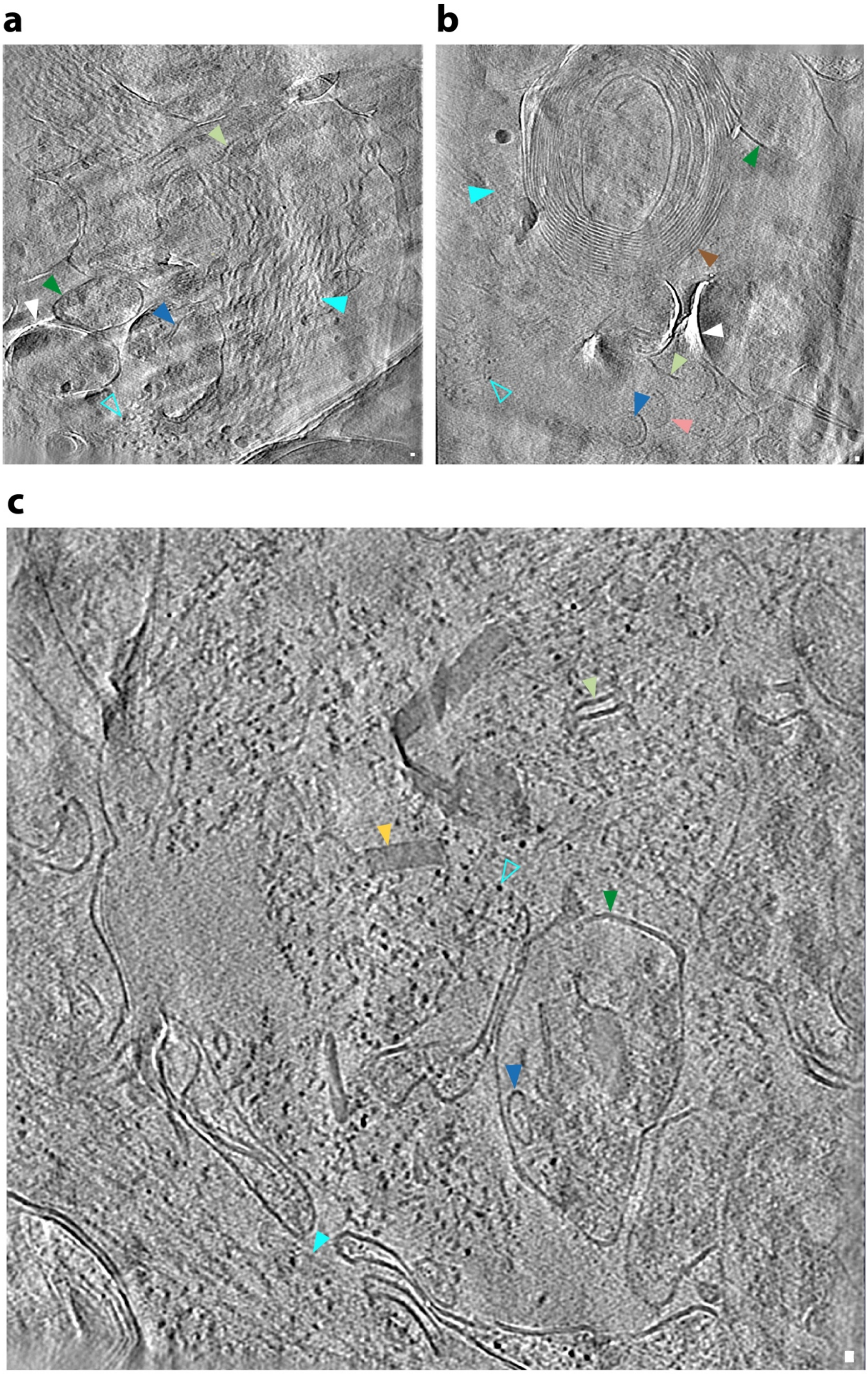


**Extended Data Figure 4. In-tissue cryoET of MX04-labelled β-amyloid plaque cryo-sections from fresh AD post-mortem brain donor.** Amyloid outside myelinated axons and subcellular compartments containing membrane-bound organelles indicates such amyloid is situated extracellularly.

**a** Tomographic slice of plaque pathology situated outside subcellular compartments in AD post-mortem brain cryo-section related to **Fig. 1g**. See also **Extended Data Movie 2.**
**b** Same as **a** but with amyloid pathology adjacent to myelinated axon.
**c** Same as **a**.
Filled and open cyan arrowhead, fibril in the *x*-*y* plane and axially (*z*-axis) of the tomogram, respectively. Brown arrowhead, myelinated axon. Dark green arrowhead, subcellular compartment. Blue arrowhead, intracellular membrane bound organelle. Light green arrowhead, open membrane sheet. Yellow arrowhead, extracellular droplet (see **Methods** '**Annotation of constituents in tomographic volumes**'). Pink arrowhead, extracellular vesicle. White arrowhead, knife damage.
Scale bar, 10 nm.

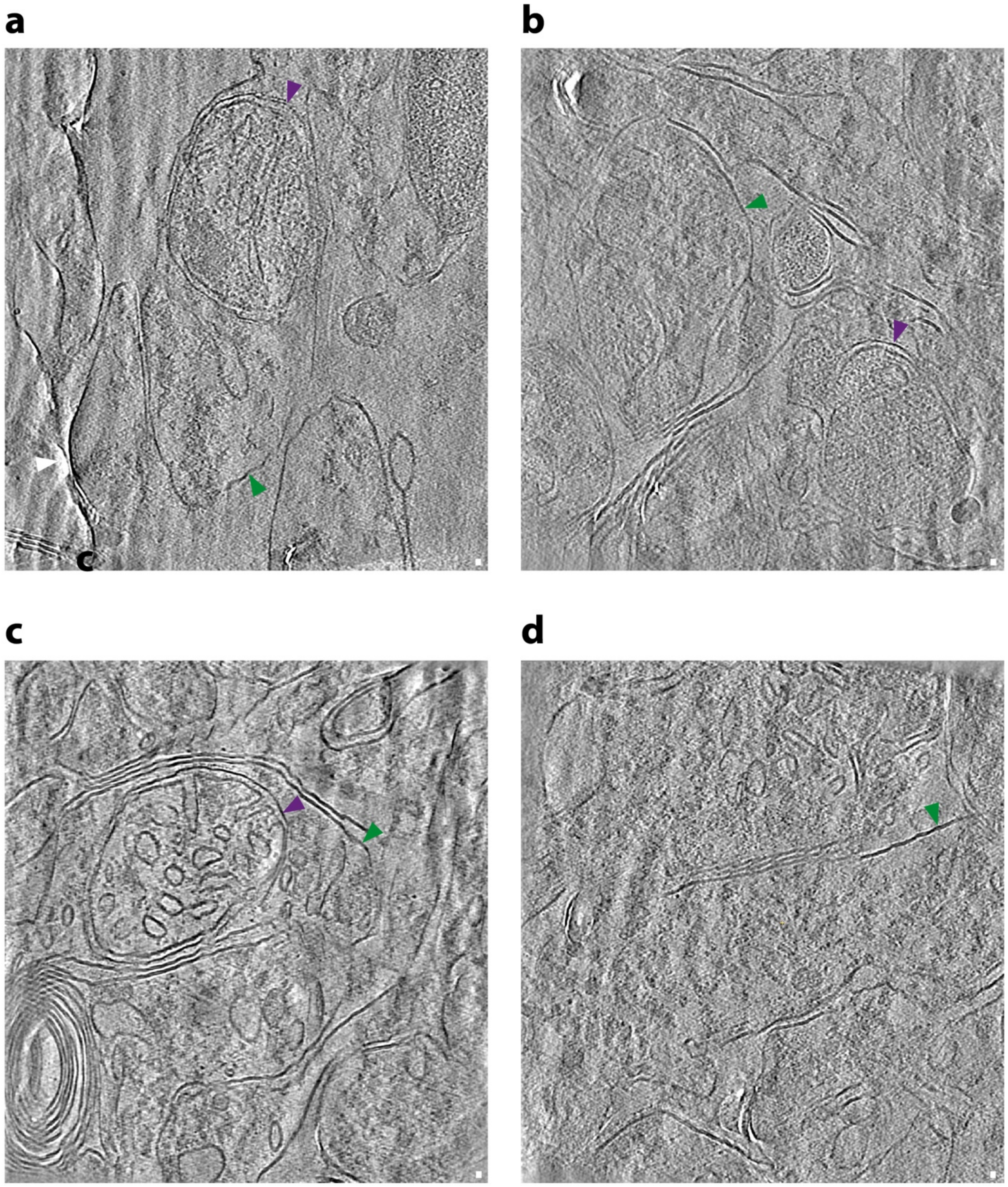


**Extended Data Figure 5. In-tissue cryoET of tissue cryo-sections from fresh non-demented control post-mortem brain donor.** Panels **a-d** show representative tomographic slices from 4 tomograms. No amyloid was observed in 64 non-demented control post-mortem donor cryo-sections (see **Extended Data Table 3**). Green arrowhead, subcellular compartment. Purple arrowhead, mitochondrion. Scale bar, 10 nm. See **Extended Data Movies 7-9**.

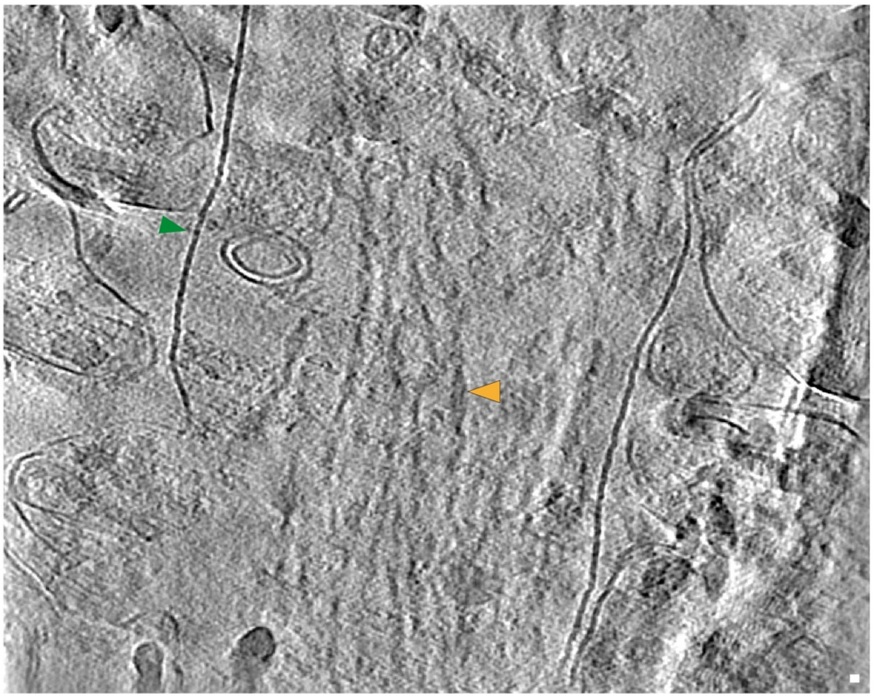

**Extended Data Figure 6. In-tissue cryoET of MX04-labelled tau inclusion cryo-sections from fresh AD post-mortem brain donor.**

Tomographic slice through dystrophic neurite. Orange arrowhead, tau filament. Green arrowhead, plasma membrane of neurite. Scale bar, 10 nm. Related to **Extended Fig. 3a** and also see **Extended Data Movie 6**.

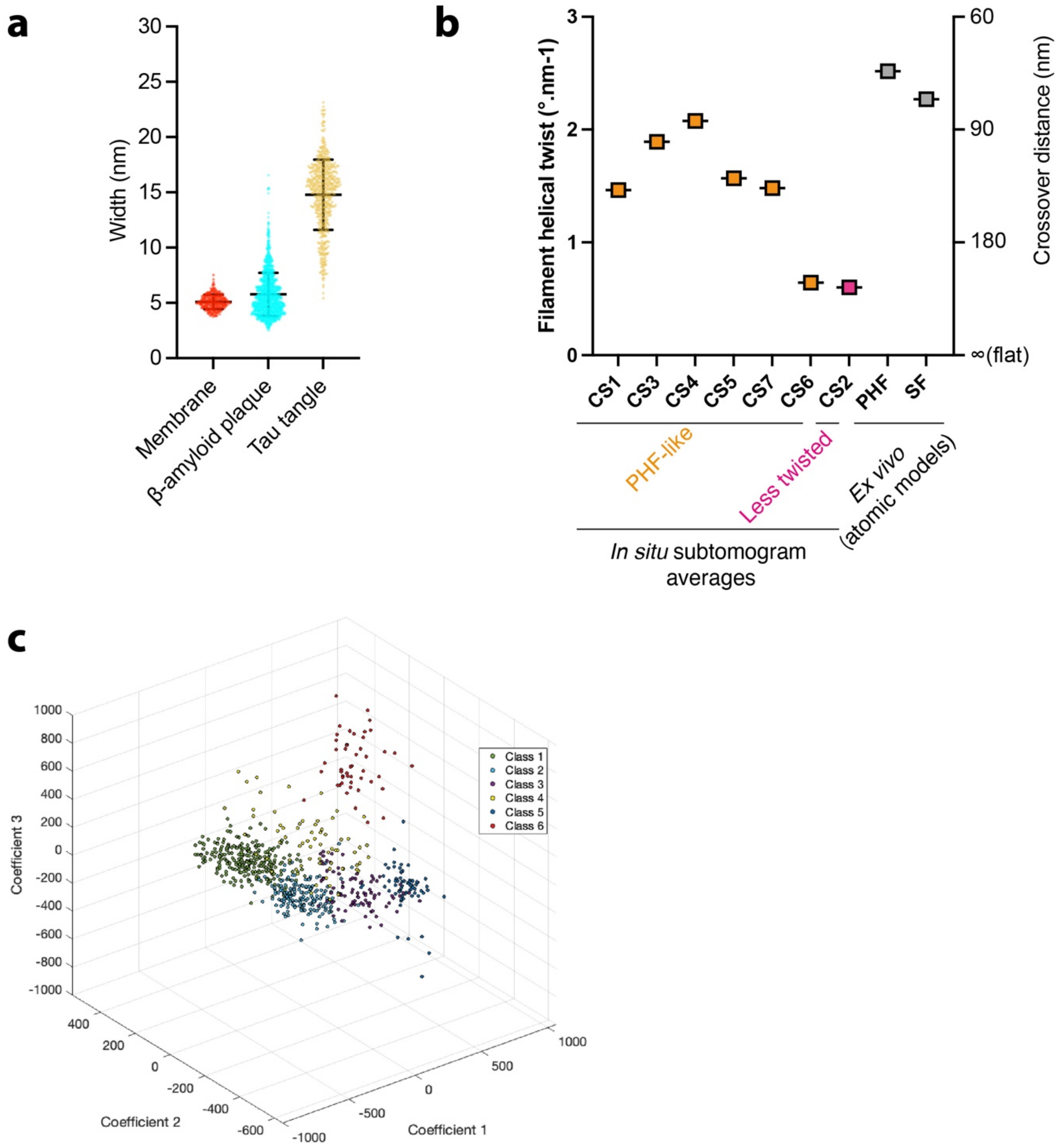


| Filament type | Tomogram ID | Class prediction from PCA analysis | | | | | | Filaments per tomogram |
|---|---|---|---|---|---|---|---|---|
| | | 1 | 2 | 3 | 4 | 5 | 6 | |
| PHF | CS1 | 0 | 53 | 12 | 8 | 2 | 0 | 75 |
| | CS3 | 0 | 118 | 3 | 0 | 0 | 0 | 121 |
| | CS4 | 0 | 0 | 0 | 0 | 58 | 0 | 58 |
| | CS5 | 253 | 0 | 1 | 3 | 0 | 0 | 257 |
| | CS7 | 0 | 0 | 0 | 0 | 2 | 48 | 50 |
| | CS6 | 0 | 0 | 6 | 57 | 0 | 0 | 63 |
| Less twisted | CS2 | 0 | 3 | 63 | 0 | 4 | 0 | 63 |
| **Total filaments in class** | | 253 | 174 | 85 | 68 | 66 | 48 | 694 |

**Extended Data Figure 7. Measurement of fibril widths in raw tomographic maps, helical twist tau subtomogram averages and unsupervised classification of tau filament subvolumes.**

**a** Scatterplot showing the width distribution of lipid membrane (used as an internal control for width measurements, n=296) and fibrils from in-tissue cryo-sections of β-amyloid plaques (n=1360) and tau tangles (n=561), respectively. Middle and top/bottom black bars indicate mean and one standard deviation.
**b** Graph showing filament helical twist (left *y*-axis) and cross-over distance (right *y*-axis) of *in situ* tau filament clusters in different locations of AD post-mortem brain from 7 cryo-section (CS1-7). Helical twist was measured using subtomogram averages from each location (see **Fig. 3i** and **Extended Data Fig. 8**). For reference the helical twist/crossover distances calculated from available *ex vivo* helically averaged atomic models of AD paired helical filaments (PHF), straight filaments (SF)[1] and chronic traumatic encephalopathy type 1 (CTE-1)[2] were also plotted.
**c** Top, 3D scatterplot showing grouping of CS1-7 tau filaments into 6 distinct classes by principal component analysis (PCA). Akaike Information Criterion (AIC) and Bayesian Information Criterion (BIC) reported highly significant positive improvement scores (3091 and 2715, respectively).
Bottom, table showing the class size, composition and identity (PHF, less twisted filaments) of tau filaments derived from subcellular locations CS1-7. Class 1 (100% CS5), class 2 (67.8% CS3, 30% CS1, >3% less-twisted CS2), class 3 (74% CS2, 14% CS1, 7% CS6, 3.5% CS3, >1% CS5), class 4 (83% CS6, 11.7% CS1, 4.4% CS5), class 5 (87% CS4, 6% CS2, 3% CS1, 3% CS7), class 6 (100% CS7).

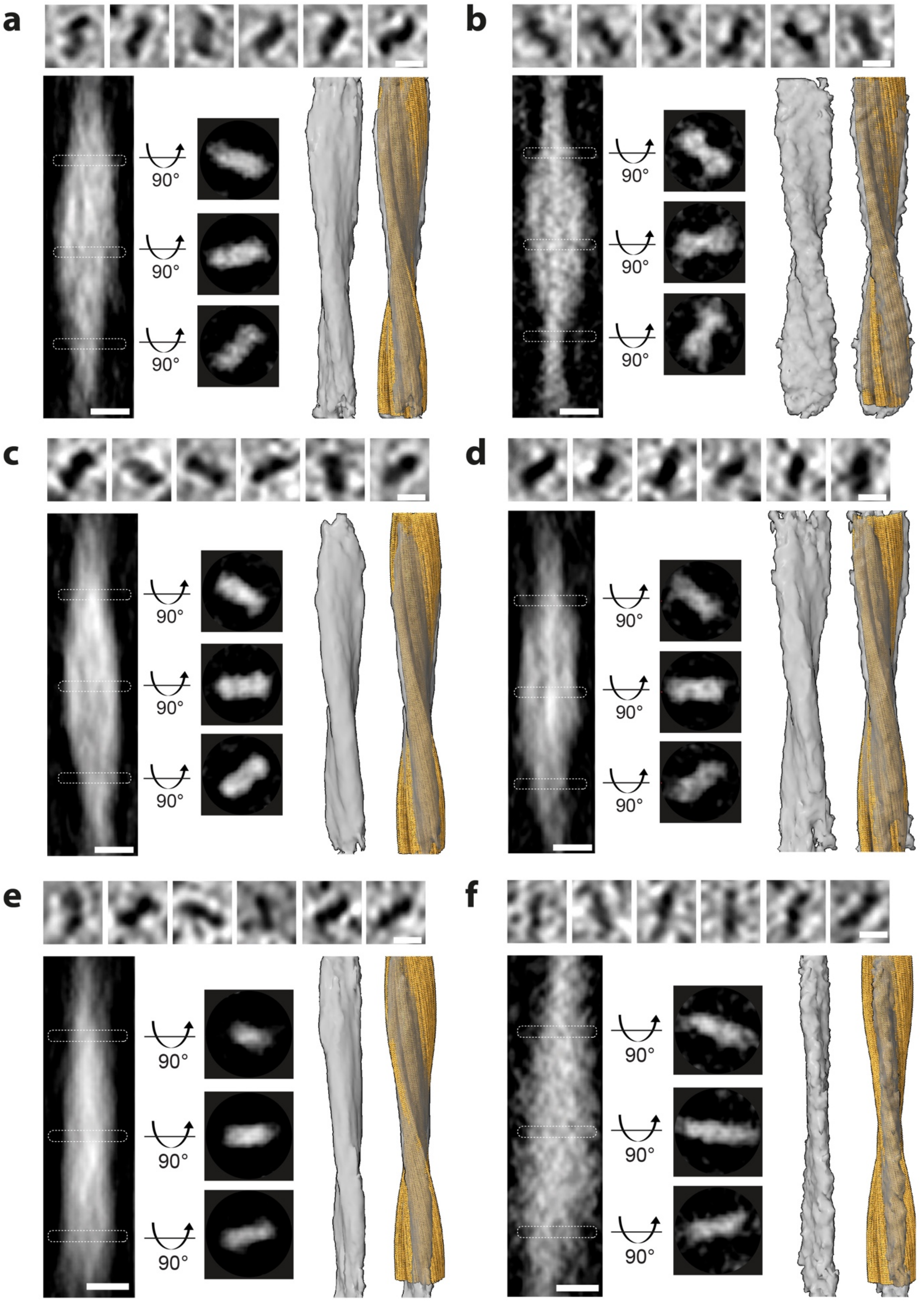


**Extended Data Figure 8. Subtomogram averaging of whole filaments (stalkInit see Methods) from tau clusters within tissue cryo-sections situated in six distinct locations, shown in panels a-e**. Top panels, representative tomographic slices of filaments picked for

subtomogram averaging. Scale bar, 10 nm. Bottom left panel, side view of tomographic slice through averaged volume showing helical twist. White dashed rectangles, the position of middle left panels along the filament axis, three top view tomographic slices at three positions (25 voxels, 23.75 nm apart). Bottom middle right and bottom right panels, subtomogram average map of tau filament with and without a helically averaged atomic model of tau PHF (yellow, PDB 5osl)[1] fitted into in the map. Scale bars, 10 nm.
Related data of a location giving the highest resolution subtomogram average is shown in **Fig. 3i-j**.
**a-e** Subtomgram averages in these five locations showed variable helical twist (see **Extended Data Fig. 7b**) and resembled PHF (see also **Extended Data Fig. 10a-e**).
**f** The subtomogram average in this location was less twisted and did not fit PHF or any other available atomic model of *ex vivo* purified tau (see **Extended Data Fig. 10f**).

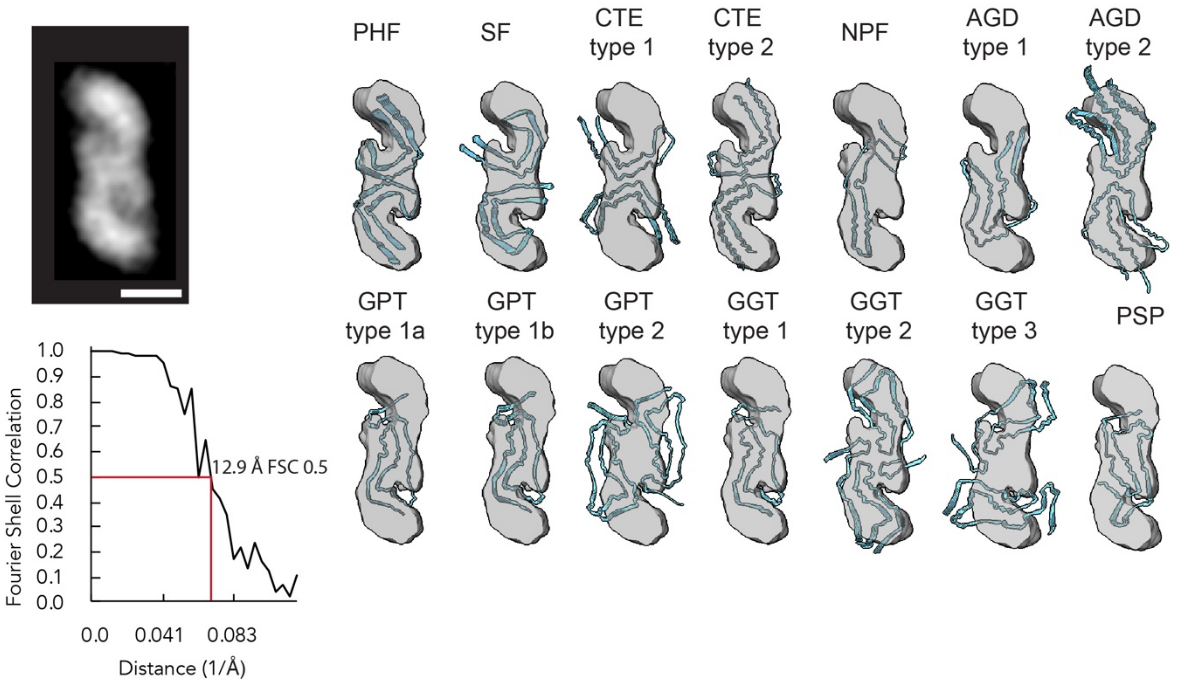


**Extended Data Figure 9. Available atomic models of tau conformers fitted into the map and resolution estimation based on Fourier shell correlation (FSC) of half-maps from subtomogram averaging of tissue cryo-section tomograms.** Related to **Fig. 3j.** Top left, slice through subtomogram averaged map. Scale bar, 10 nm. Bottom left, graph showing spatial frequency (1/Å versus FSC. Right, panel of tau conformers fitted into subtomogram average map: PHF, AD paired helical filaments (PDB: 5O3L). SF, AD straight filaments (PDB 5O3T)[1]. CTE type 2 and 2, Chronic traumatic encephalopathy types 1 and 2 (PDB: 6NWP and 6NWQ), respectively[2]. NPF, narrow picks filament (PDB 6GX5)[3]. AGD type 1 and 2, Argyrophilic grain disease types 1 and 2 (PDB 7P6D and 7P6E). GPT, GGT-PSP type 1a, 1b and 2, (PDB 7P6A, 7P6B, 7P6C), respectively. GGT, globular glial tauopathy types 1, 2 and 3 (PDB 7P66, 7P67, 7P68), respectively. PSP, progressive supranuclear palsy (PDB 7P65)[4]. The PHF is the only known tau amyloid structure that fits the map.

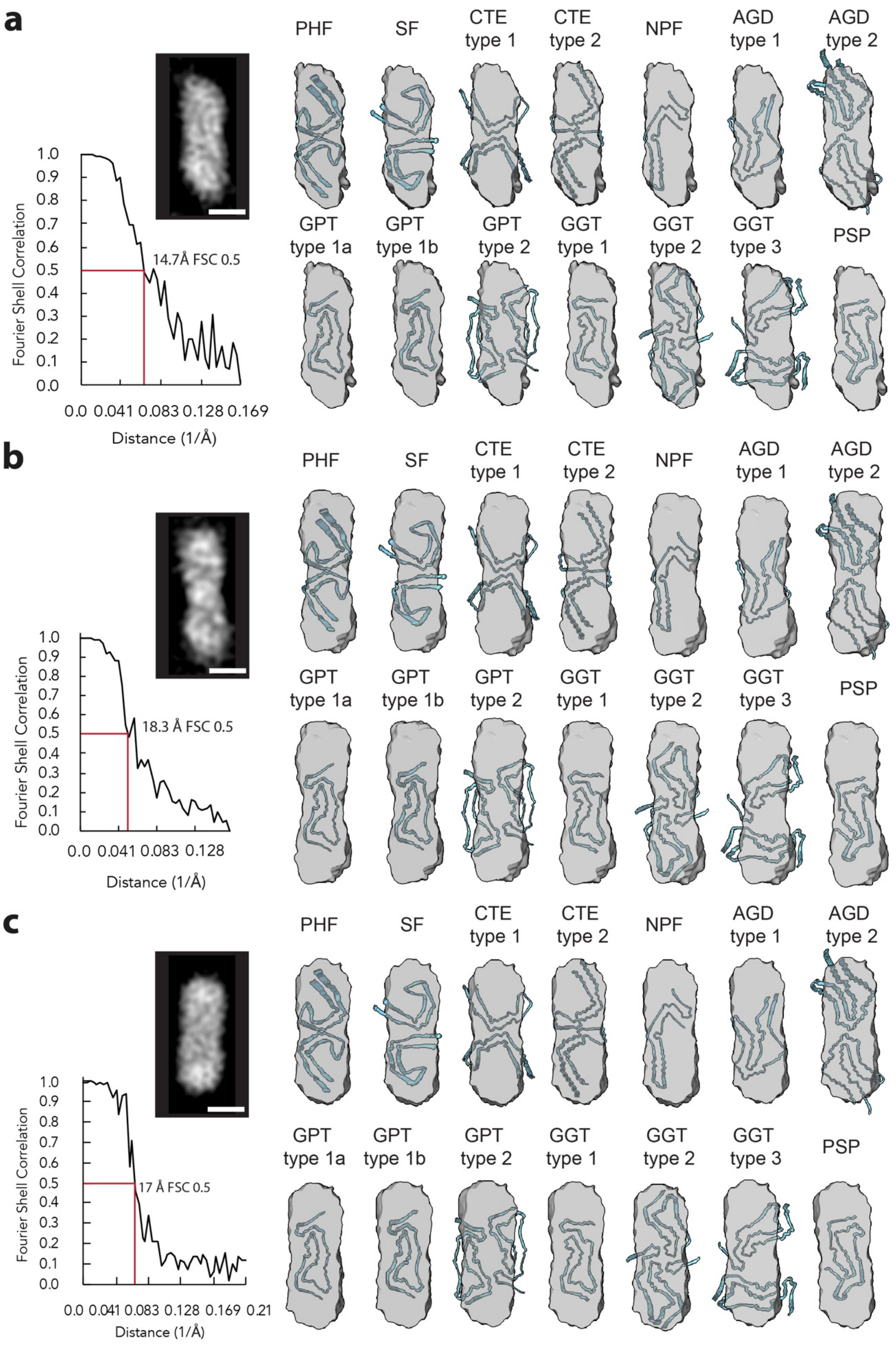

a
PHF
SF
CTE type 1
CTE type 2
NPF
AGD type 1
AGD type 2
GPT type 1a
GPT type 1b
GPT type 2
GGT type 1
GGT type 2
GGT type 3
PSP
Fourier Shell Correlation
1.0
0.9
0.8
0.7
0.6
0.5
0.4
0.3
0.2
0.1
0.0
14.7Å FSC 0.5
0.0 0.041 0.083 0.128 0.169
Distance (1/Å)
b
PHF
SF
CTE type 1
CTE type 2
NPF
AGD type 1
AGD type 2
GPT type 1a
GPT type 1b
GPT type 2
GGT type 1
GGT type 2
GGT type 3
PSP
Fourier Shell Correlation
1.0
0.9
0.8
0.7
0.6
0.5
0.4
0.3
0.2
0.1
0.0
18.3 Å FSC 0.5
0.0 0.041 0.083 0.128
Distance (1/Å)
c
PHF
SF
CTE type 1
CTE type 2
NPF
AGD type 1
AGD type 2
GPT type 1a
GPT type 1b
GPT type 2
GGT type 1
GGT type 2
GGT type 3
PSP
Fourier Shell Correlation
1.0
0.9
0.8
0.7
0.6
0.5
0.4
0.3
0.2
0.1
0.0
17 Å FSC 0.5
0.0 0.041 0.083 0.128 0.169 0.21
Distance (1/Å)

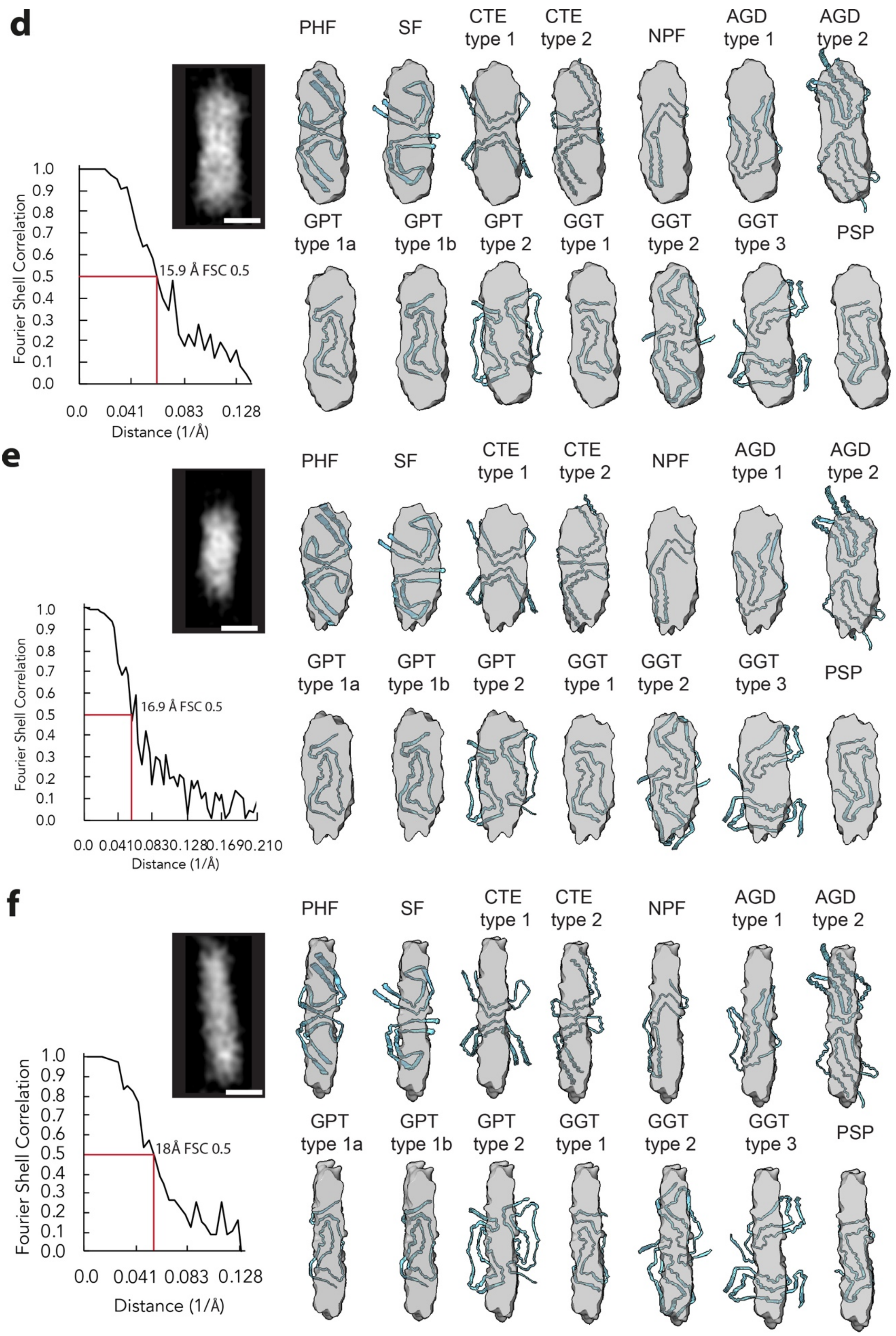


**Extended Data Figure 10. Available atomic models of known tau amyloid conformers fitted into the map and resolution estimation based on Fourier shell correlation (FSC) of half-maps from subtomogram averaging of different tau filament clusters a-f**. Top left, slice through subtomogram averaged map. Scale bar, 10 nm. Bottom left, graph showing spatial frequency

(1/Å) versus FSC. Right, panel of tau conformers fitted into subtomogram average map: PHF, AD paired helical filaments (PDB: 5O3L). SF, AD straight filaments (PDB 5O3T)[1]. CTE type 1 and 2, Chronic traumatic encephalopathy types 1 and 2 (PDB: 6NWP and 6NWQ), respectively[2]. NPF, narrow picks filament (PDB 6GX5)[3]. AGD type 1 and 2, Argyrophilic grain disease types 1 and 2 (PDB 7P6D and 7P6E). GPT, GGT-PSP type 1a, 1b and 2, (PDB 7P6A, 7P6B, 7P6C), respectively. GGT, globular glial tauopathy types 1, 2 and 3 (PDB 7P66, 7P67, 7P68), respectively. PSP, progressive supranuclear palsy (PDB 7P65)[4].
**a-e** Subtomogram averages from these locations resembled paired helical filaments.
**f** Subtomogram average from this location did not fit any available atomic model of *ex vivo* purified tau amyloid.

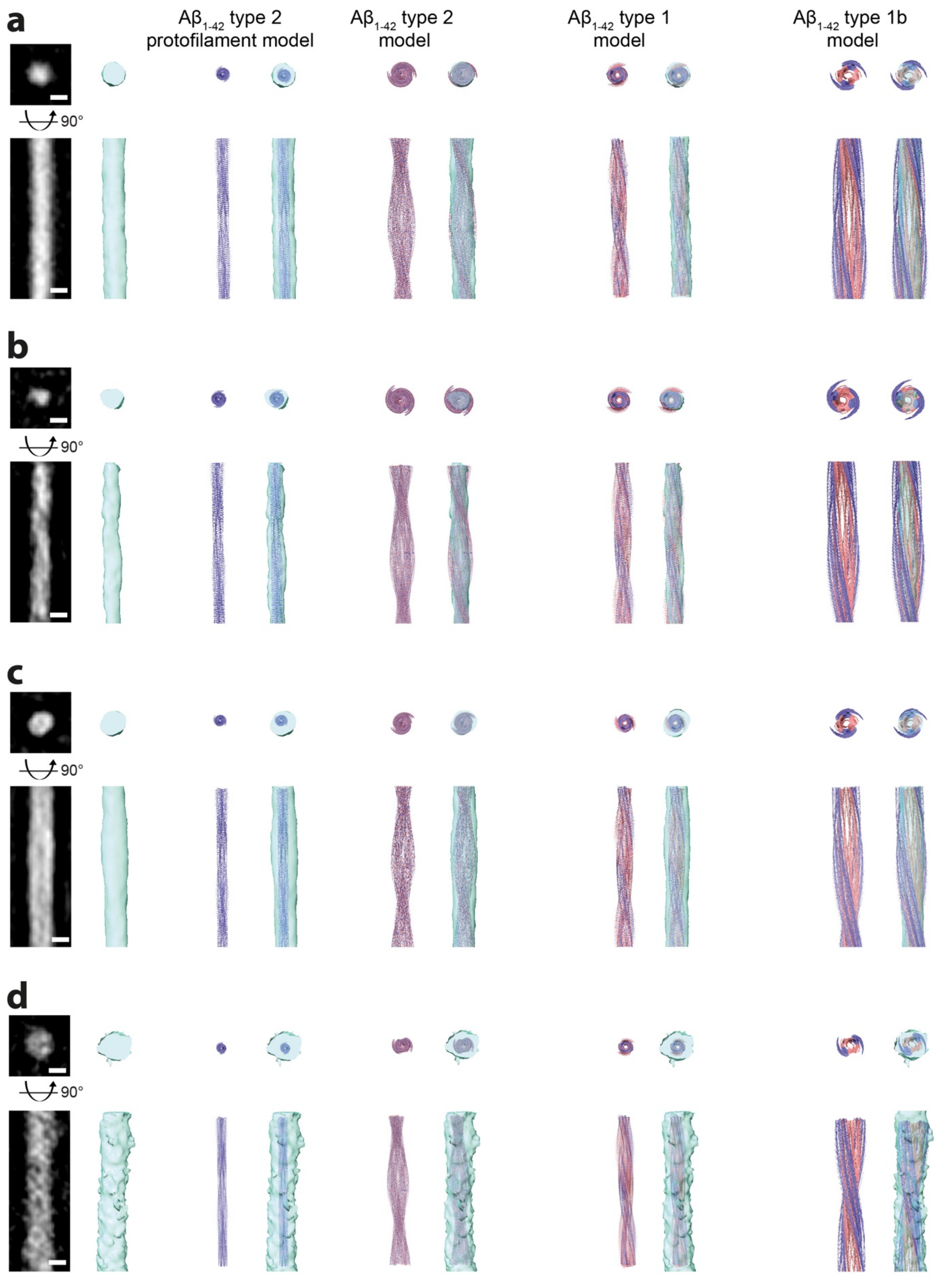


**Extended Data Figure 11. Subtomogram averaging of Aβ fibrils from MX04-labelled β-amyloid plaque cryo-sections.** Pairs of panels showing, top and bottom, top and side view of fibril, respectively.

Left pair, tomographic slice and map of subtomogram average. Scale bar, 5 nm.

Middle left pair, model of A$\beta_{1\text{-}42}$ type 2 (PDB 7Q4M)[5] protofilament alone and fitted into subtomogram average map.
Middle pair, atomic model of A$\beta_{1\text{-}42\ \text{type}}$ 2 (PDB 7Q4M)[5] alone and fitted into subtomogram average map.
Middle right pair, atomic model of A$\beta_{1\text{-}42}$ type 1 (PDB 7Q4B)[5] alone and fitted into subtomogram average map.
Right pair, atomic model of A$\beta_{1\text{-}42}$ type 1b (composed of two A$\beta_{1\text{-}42}$ type 1 fibrils PDB 7Q4B)[5] alone and fitted into subtomogram average map.
**a** Subtomogram averaging of 100 fibrils (of all widths) from one tomogram.
**b** Subtomogram averaging of 20 protofilament-like rods from one tomogram.
**c** Subtomogram averaging of 42 fibrils from one tomogram.
**d** Subtomogram averaging of 42 thick fibrils from a different tomogram.

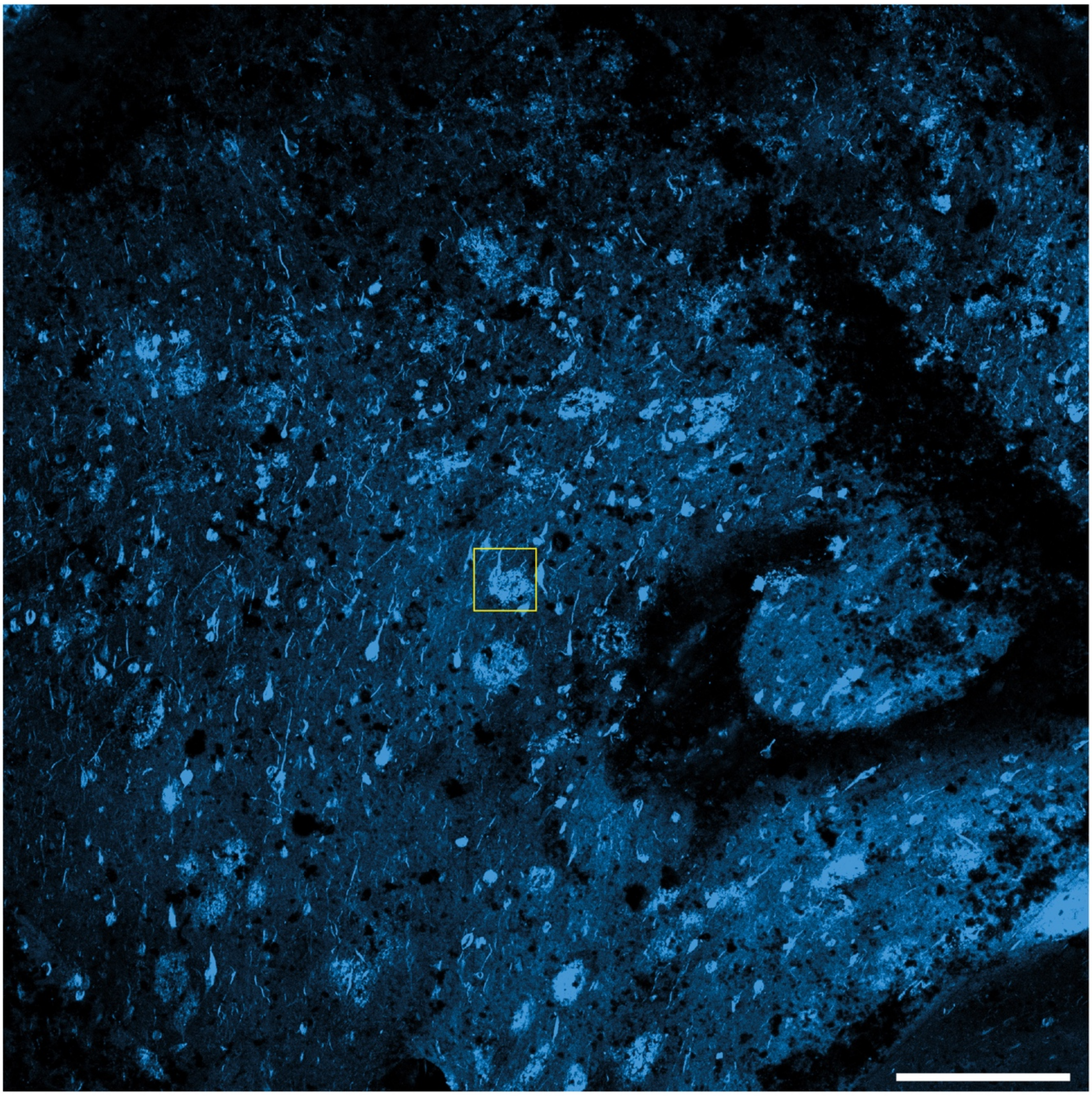

**Extended Data Figure 12. Confocal cryoFM of MX04-labelled post-mortem AD brain.** Overview confocal fluorescence microscopy image (maximum intensity projection of fresh, MX04-labelled, high-pressure frozen human postmortem tissue sample, 10x/0.4NA objective as described in the **Methods** Yellow square, region targeted for cryoFIB-SEM liftout (see **Fig. 4** and **Extended Data Fig. 13**). Scale bar, 200 µm.

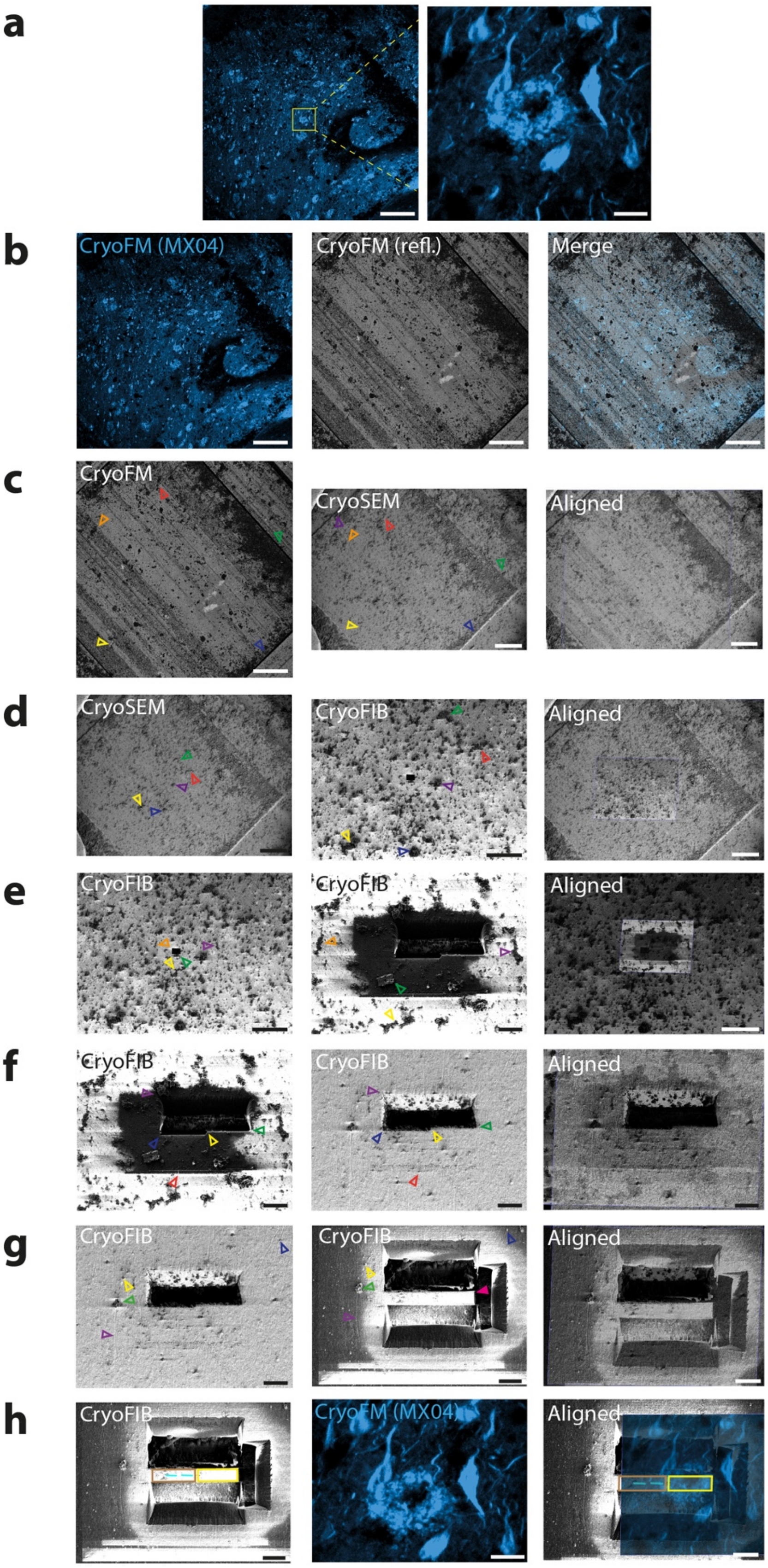


**Extended Data Figure 13. Correlated cryoFM-FIB-SEM liftout targeting of MX04-labelled tau pathology in high-pressure frozen post-mortem AD brain.**

**a** CryoFM showing MX04-labelled amyloid pathology targeted for cryoFIB-SEM liftout . Left, 10x objective overview image. Scale bar, 200 µm. Right, 100x objective confocal image. Scale bar, 20 µm. CryoFM images were aligned in ZEN. See also **Extended Data Figure 12**.
**b** CryoFM overview image (5x magnification) of MX04-labelled high-pressure frozen post-mortem AD brain. Left, reflection mode image showing surface information. Middle, MX04 cryoFM showing amyloid. Right, Merged image of left and middle. Scale bar, 200 µm.
**c-h** Alignment of cryoFM, FIB and SEM images to target liftout in ZEN Connect. Images on left were aligned with middle image using fiducials indicated by open arrowheads.
**c** Alignment between cryoSEM normal overview and cryoSEM normal high-magnification image of MX04-labelled liftout target. Left, cryoSEM normal overview. Middle, cryoSEM normal high-magnification image. Right, merged image showing alignment. Scale bar, 200 µm.
**d** Alignment between cryoSEM normal high-magnification and cryoFIB normal image. Left, cryoSEM normal high magnification. Scale bar, 200 µm. Middle, cryoFIB normal image. Scale bar, 100 µm. Right, merged image showing alignment. Scale bar, 200 µm.
**e** Alignment between cryoFIB normal image and cryoFIB high-magnification image before and after milling a trench near target for liftout, used for fiducial alignment in **g.** Left and middle, cryoFIB high-magnification before and after cryoFIB milling trench (scale bar, 100 µm and 20 µm), respectively. Right, merged image showing alignment. Scale bar, 20 µm.
**f** Alignment between cryoFIB images before and after surface cleaning, sputtercoating and cold deposition of platinum precursor. Left, cryoFIB high-magnification image. Middle, cryoFIB high-magnification image following cold deposition of GIS. Right, merged image showing alignment. Scale bar, 20 µm.
**g** Alignment of cryoFIB images before and after trenches were milled in front, behind and to the right side to prepare tissue chunk for targeted liftout (see **Extended Fig. 14**). Left and middle, cryoFIB high-magnification image before and after cryoFIB milling preparation of tissue chunk. Magenta arrowhead, cryoFIB-milled tissue chunk. Right, merged image showing alignment. Scales bar, 20 µm.
**h** Images showing the result of alignments (from **a** to **h**) to target MX04-labelled tau for cryoFIB-SEM liftout. Left, cryoFIB image of tissue chunk prepared for liftout. Middle, cryoFM 100x magnification confocal image of MX04. Right, Right, merged image showing alignment. Brown and yellow rectangles correspond to left and right regions of serial chunk liftout, respectively. Cyan lines, locations of lamella windows (see **Extended Data Figure 14**). The right chunk was lost during transfer from cryoFIB to Krios TEM.

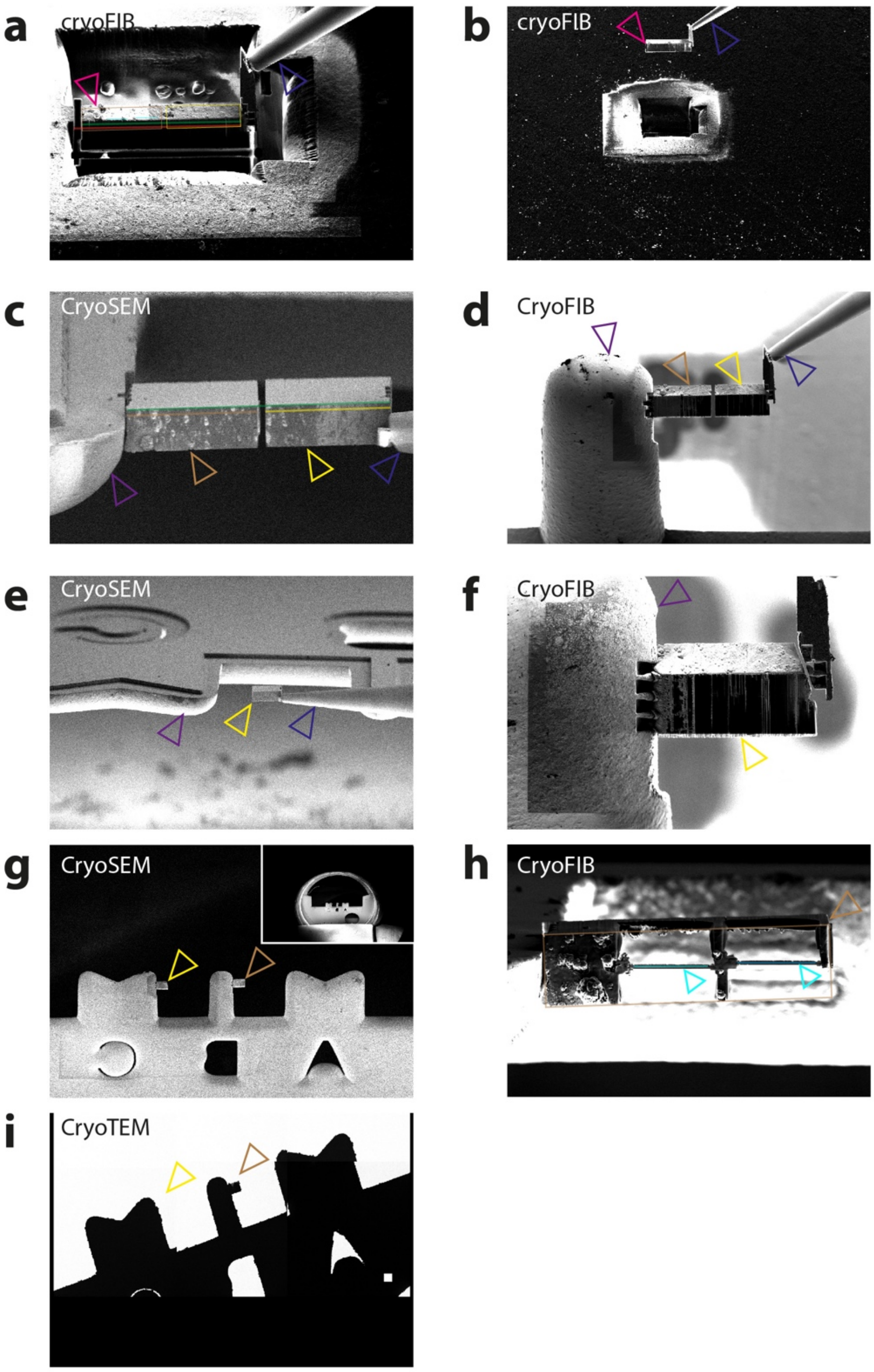


**Extended Data Figure 14. CryoFIB-SEM liftout of MX04-labelled post-mortem AD brain containing tau.** Lines and rectangles tracing the tissue chunk during cryoFIB cuts and thinning were used for cryoCLEM (see **Extended Data Fig. 13i**). Red semi-transparent line, length of tissue chunk before liftout. Green semi-transparent line, length of tissue chunk during liftout. Brown rectangle, left tissue chunk. Yellow rectangle, right tissue chunk (lost during sample transfer from cryoFIB-SEM to cryoEM). Cyan lines, region targeted for cryoFIB milling lamella windows within tissue chunk.

**a** CryoFIB image of tissue chunk after final left side cut to detach tissue chunk. Open magenta arrowhead, detached tissue chunk. Open blue arrowhead, liftout tool (copper block linking micromanipulator needle) attached to right side of tissue chunk.

**b** CryoFIB image showing tissue chunk liftout. Magenta arrowhead, tissue chunk. Blue open arrowhead, liftout tool.
**c** CryoSEM images showing attachment of the left tissue chunk to EM grid and cryoFIB cut between the left and right tissue chunks. Purple open arrowhead, EM grid. Yellow open arrowhead, left tissue chunk. Green open arrowhead, right tissue chunk. Blue open arrowhead, liftout tool.
**d** Same as **c,** but cryoFIB image.
**e** CryoSEM image showing left tissue chunk. Arrowheads same as **c**.
**f** CryoFIB showing right tissue chunk after attachment to second location on EM grid and the final cryoFIB cut to detach liftout tool. Arrow heads same as **c**.
**g** CryoSEM showing serial attachment of the left and right tissue chunks. Green and yellow open arrowheads, left and right tissue chunks, respectively. Inset, overview image showing mounted autogrid with clipped halfmood Omniprove EM grid.
**h** CryoFIB normal view image (56° stage tilt) of the left chunk after cryoFIB thinning to produce two 130-200 nm thick, ~8 µm wide, ~15 µm deep lamellae windows. Cyan open arrowhead, lamella. Brown open arrowhead, tissue chunk. Brown rectangle, tissue chunk dimensions (used for cryoCLEM see **Extended Data Fig. 13h**).
**i** CryoEM overview showing left and right cryoFIB-milled tissue chunk attachment positions, left was lost and right remained during transfer to cryoEM, respectively. Arrowheads, same as **g**.

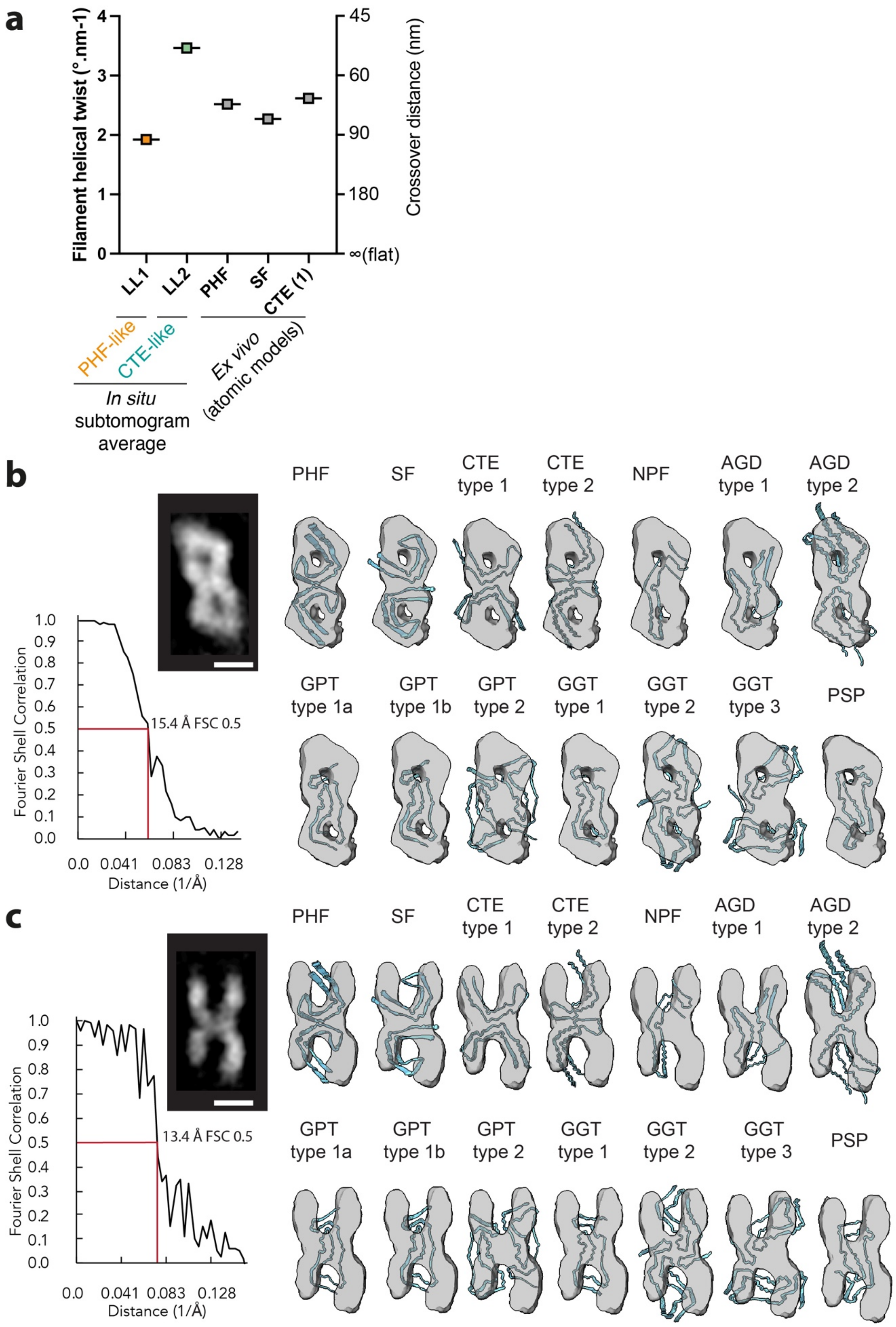


**Extended Data Figure 15. Distinct structures of filaments from different tau clusters in cryoFIB-SEM liftout lamella tomograms.**

**a** Graph showing filament helical twist (left *y*-axis) and cross-over distance (right *y*-axis) of *in situ* tau filament clusters in different locations of AD post-mortem brain from 2 cryoFIB-SEM liftout lamella tomograms (LL1-2). Helical twist was measured using subtomogram averages from each location (see **Fig. 4h and Fig. 4j)**. For reference the helical twist/crossover distances calculated from available *ex vivo* helically averaged atomic models of AD paired helical filaments (PHF), straight filaments (SF)[1] and chronic traumatic encephalopathy type 1 (CTE-1)[2] were also plotted.

**b and c** Available atomic models of tau amyloid conformers fitted into the map and resolution estimation based on Fourier shell correlation (FSC) of half-maps from subtomogram averaging of two different tau clusters in cryoFIB-SEM liftout tomograms.

Top left, slice through subtomogram averaged map. Bottom left, graph showing spatial frequency (1/Å) versus FSC. Right, panel of known tau amyloid conformers fitted into subtomogram average map: PHF, AD paired helical filaments (PDB: 5O3L). SF, AD straight filaments (PDB 5O3T)[1]. CTE type 1 and 2, Chronic traumatic encephalopathy types 1 and 2 (PDB: 6NWP and 6NWQ), respectively[2]. NPF, narrow picks filament (PDB 6GX5)[3]. AGD type 1 and 2, Argyrophilic grain disease types 1 and 2 (PDB 7P6D and 7P6E). GPT, GGT-PSP type 1a, 1b and 2, (PDB 7P6A, 7P6B, 7P6C), respectively. GGT, globular glial tauopathy types 1, 2 and 3 (PDB 7P66, 7P67, 7P68), respectively. PSP, progressive supranuclear palsy (PDB 7P65)[4].

**a** Related to **Fig, 4i**. Subtomogram average of paired helical filaments in post-mortem AD tau thread prepared by cryoFIB-SEM liftout.

**b** Related to **Fig, 4k**. Subtomogram average of CTE-like filaments in post-mortem AD brain prepared by cryoFIB-SEM liftout.

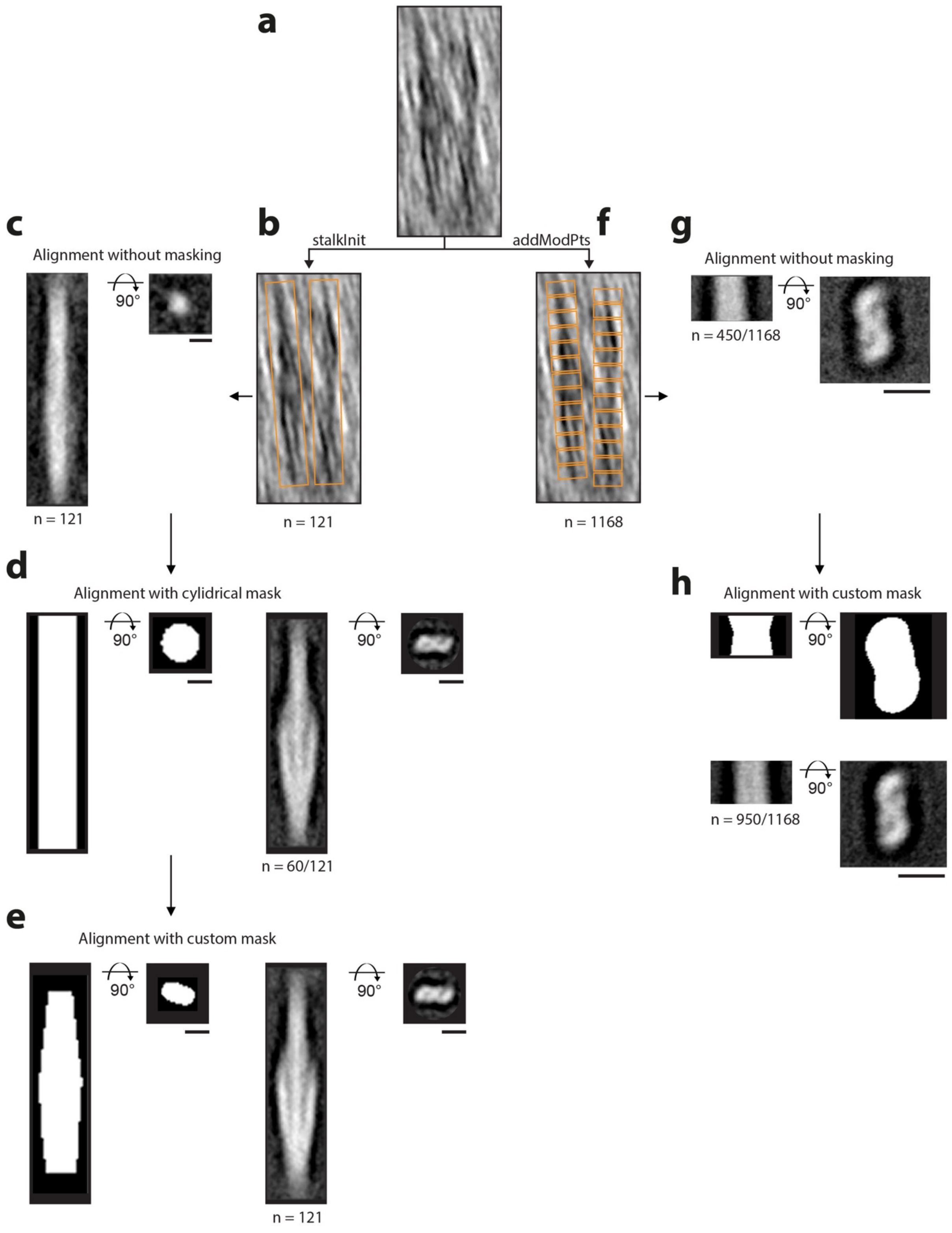


**Extended Data Figure 16. Subtomogram averaging scheme of tau pathology in tissue cryo-sections. Related to Fig. 3i-j.**

**a** Tomographic slice showing side (*z-y* plane) view of two representative filaments in raw tomographic volume.

**b** For each picked filament, model points were placed at opposite poles of the filament to generate a two-point contour running along the filament axis. The PEET stalkInit command was used to generate single-point model files (coordinates of the head, tail and centroid of each picked filament subvolume), initial motive list (MOTL) and rotation axis files. These files were used for the initial subvolume alignment and averaging. Orange rectangles, filament subvolume.

**c** Left and right panel, tomographic slice side and top view of averaged subvolumes (n = 121) produced from the first round of alignments without masking, respectively.

**d** Subsequent rounds of subvolume alignment and averaging were performed with updated motive lists, rotation axes and model point files, generated using createAlignedModel. A cylindrical mask with blurred edges (4 voxels) was applied to the reference for subvolume alignment. Left and left-middle panel, side and top view slice of cylindrical mask used for subvolume alignment, respectively. Right-middle and right panel, tomographic slice showing side and top view of the subtomogram average produced from cylindrical masked alignment of 60 filament subvolumes, respectively.

**e** In the next round of subvolume alignment and averaging, createAlignedModel was used again, to generate updated motive lists, rotation axes and model point files (see methods). A custom mask with blurred edges (4 voxels) was applied to the reference for subvolume alignments. Left and left-middle panel, side and top view slice of custom mask used for subvolume alignment. Right-middle and right panel, side and top view tomographic slice of the subtomogram average produced using custom masking for the alignment of each filament subvolume, respectively.

**f** Representative tau filaments picked in 40 voxel increments (orange rectangles) using the addModPts command in PEET. The addModPts generated model files were used for the initial subvolume alignment and averaging.

**g** Left and right panel, tomographic slice showing side and top view of a portion of averaged subvolumes generated by addModPts (n = 450/1168) without masking, respectively.

**h** Subsequent rounds of subvolume alignment and averaging were performed with updated motive lists, rotation axes and model point files (see methods), generated using createAlignedModel. A custom mask with blurred edges (5 voxels) was applied to the reference for subvolume alignment. Top left and right panel, side and top view slice of the custom mask used for subvolume alignment. Bottom left and right panel, tomographic slice showing side and top view of the subtomogram average produced using custom masking for the alignment of 950 filament subvolumes.

Scale bar, 10 nm.

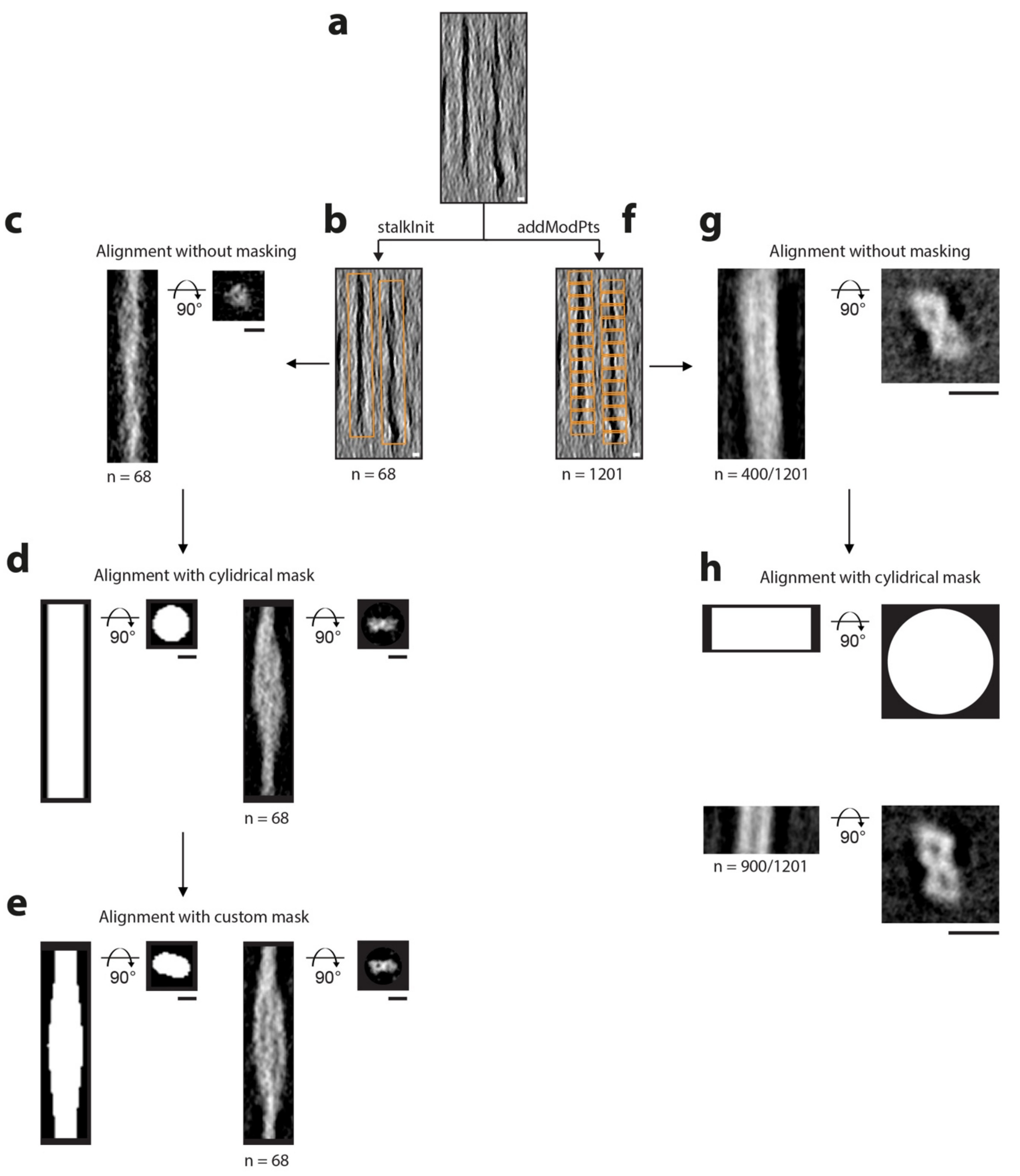


**Extended Data Figure 17. Subtomogram averaging scheme of tau pathology in tissue cryoFIB-SEM liftout lamella. Related to Fig. 4h-i.**

**a** Tomographic slice showing side (*z*-*y* plane) view of two representative filaments in raw tomographic volume.

**b** For each picked filament, model points were placed at opposite poles of the filament to generate a two-point contour running along the filament axis. The PEET stalkInit command was used to generate single-point model files (coordinates of the head, tail and centroid of each picked filament subvolume), initial motive list (MOTL) and rotation axis files. These files were used for the initial subvolume alignment and averaging. Orange rectangles, filament subvolume.

**c** Left and right panel, tomographic slice side and top view of averaged subvolumes (n = 68) produced from the first round of alignments without masking, respectively.

**d** Subsequent rounds of subvolume alignment and averaging were performed with updated motive lists, rotation axes and model point files, generated using createAlignedModel. A cylindrical mask with blurred edges (4 voxels) was applied to the reference for subvolume alignment. Left and left-middle panel, side and top view slice of cylindrical mask used for subvolume alignment, respectively. Right-middle and right panel, tomographic slice showing side and top view of subtomogram average produced from cylindrical masked alignment of 68 filament subvolumes, respectively.
**e** In the next round of subvolume alingment and averaging, createAlignedModel was used again, to generate updated motive lists, rotation axes and model point files (see methods). A custom mask with blurred edges (4 voxels) was applied to the reference for subvolume alignments. Left panel, slice of custom mask used for subvolume alignment. Right-middle and right panel, tomographic slice showing side and top view of subtomogram average produced using custom masking for the alignment of each filament subvolume, respectively.
**f** Representative tau filaments picked in 40 voxel increments (orange rectangles) using the addModPts command in PEET. The addModPts generated model files were used for the initial subvolume alignment and averaging.
**g** Left and right panel, tomographic slice showing side and top view of a portion of averaged subvolumes generated by addModPts (n = 400/1201) without masking, respectively.
**h** Subsequent rounds of subvolume alignment and averaging were performed with updated motive lists, rotation axes and model point files (see methods), generated using createAlignedModel. A cylindrical mask with blurred edges (5 voxels) was applied to the reference for subvolume alignment. Top left and right panel, side and top view slice of the cylindrical mask used for subvolume alignment. Bottom left and right panel, tomographic slice showing side and top view of the subtomogram average produced using cylindrical masking for the alignment of (900/1201) filament subvolumes, respectively.
Scale bar, 10 nm.

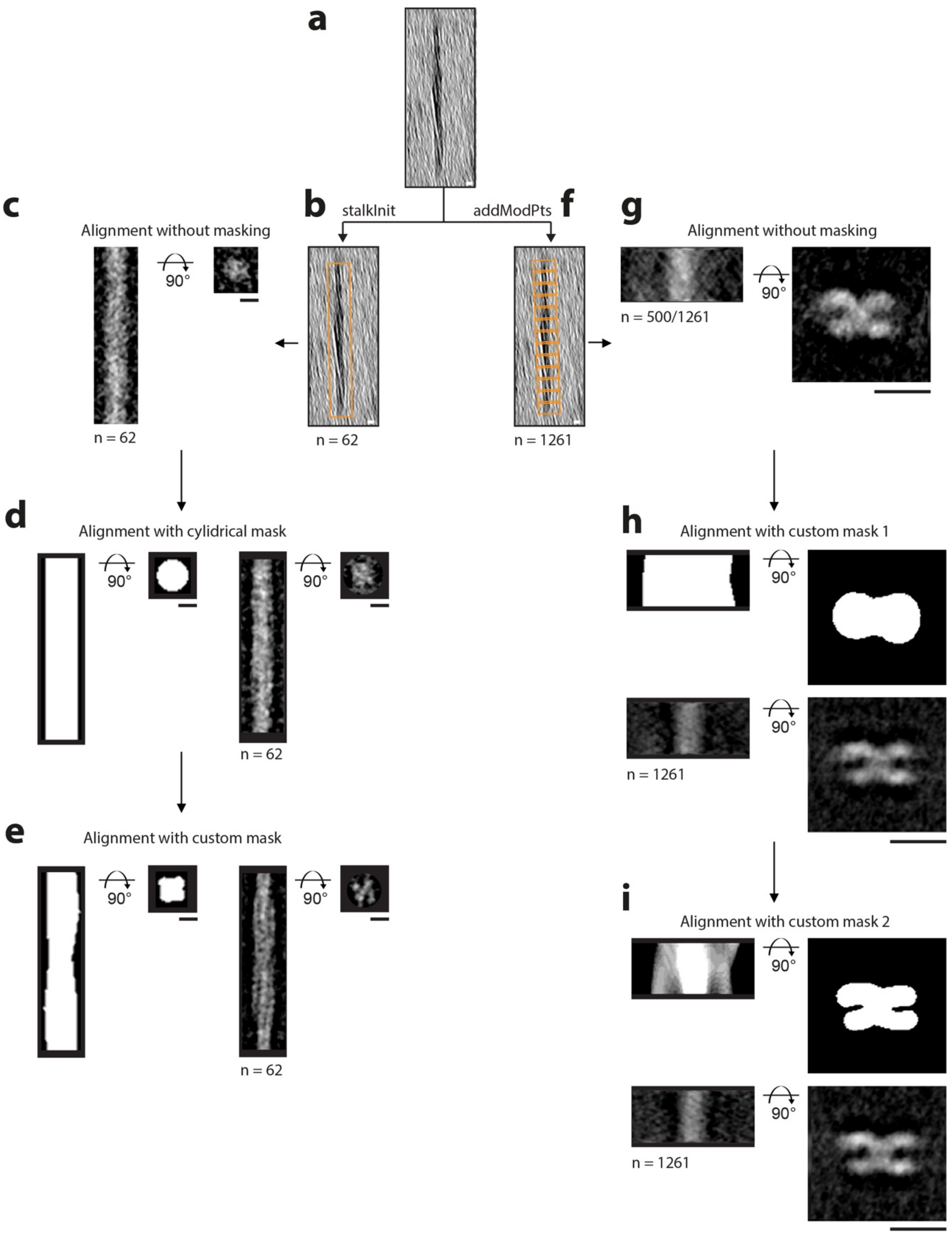


**Extended Data Figure 18. Subtomogram averaging scheme of MX04-labelled tau pathology in tissue cryoFIB-SEM liftout lamella. Related to Fig. 4j-k.**

**a** Tomographic slice showing side (*z-y* plane) view of two representative filaments in raw tomographic volume.

**b** For each picked filament, model points were placed at opposite poles of the filament to generate a two-point contour running along the filament axis. The PEET stalkInit command was used to generate single-point model files (coordinates of the head, tail and centroid of each picked filament subvolume), initial motive list (MOTL) and rotation axis files. These files

were used for the initial subvolume alignment and averaging. Orange rectangles, filament subvolume.
**c** Left and right panel, tomographic slice side and top view of averaged subvolumes (n = 62) produced from the first round of alignments without masking, respectively.
**d** Left and left-middle panel, side and top view slice of cylindrical mask used for subvolume alignment, respectively. Right-middle and right panel, side and top view tomographic slice of the subtomogram average produced from cylindrical masked alignment of 62 filament subvolumes, respectively.
**e** Subsequent rounds of subvolume alignment and averaging were performed with updated motive lists, rotation axes and model point files, generated using createAlignedModel. A custom mask with blurred edges (4 voxels) was applied to the reference for subvolume alignments. Left panel, slice of custom binary mask used for subvolume alignment. Right-middle and right panel, tomographic slice showing side and top view of subtomogram average produced using custom masking for the alignment of each filament subvolume, respectively.
**f** Representative tau filaments picked in 40 voxel increments (orange rectangles) using the addModPts command in PEET. The addModPts generated model files were used for the initial subvolume alignment and averaging.
**g** Left and right panel, tomographic slice showing side and top view of a portion of averaged subvolumes generated by addModPts (n = 500/1261) without masking, respectively.
**h** Subsequent rounds of subvolume alignment and averaging were performed with updated motive lists, rotation axes and model point files (see methods), generated using createAlignedModel. A custom mask with blurred edges (5 voxels) was applied to the reference for subvolume alignment. Top left and right panel, side and top view slice of custom mask used for subvolume alignment. Bottom left and right panel, tomographic slice showing side and top view of the subtomogram average produced using custom masking for the alignment of 1261 filament subvolumes.
i Final rounds of subvolume alignment and averaging were performed as in **h** but with a new custom mask.
Scale bar, 10 nm.

### Extended Data Figure References
